# Substrate stiffness and shear stress collectively regulate the inflammatory phenotype in cultured human brain microvascular endothelial cells

**DOI:** 10.1101/2025.03.31.646228

**Authors:** Alexis Yates, Heather Murray, Andrew Kjar, Daniel Chavarria, Haley Masters, Hyosung Kim, Alexander P. Ligocki, Angela L. Jefferson, Ethan S. Lippmann

## Abstract

Brain endothelial cells experience mechanical forces in the form of blood flow-mediated shear stress and underlying matrix stiffness, but intersectional contributions of these factors towards blood-brain barrier (BBB) impairment and neurovascular dysfunction have not been extensively studied. Here, we developed *in vitro* models to examine the sensitivity of primary human brain microvascular endothelial cells (BMECs) to substrate stiffness, with or without exposure to fluid shear stress. Using a combination of molecular profiling techniques, we show that BMECs exhibit an inflammatory signature at both the mRNA and protein level when cultured on gelatin substrates of intermediate stiffness (∼30 kPa) versus soft substrates (∼6 kPa). Exposure to modest fluid shear stress (1.7 dyne/cm^2^) partially attenuated this signature, including reductions in levels of soluble chemoattractants and surface ICAM-1. Overall, our results indicate that increased substrate stiffness promotes an inflammatory phenotype in BMECs that is dampened in the presence of fluid shear stress.

## 2. Background

The blood-brain barrier (BBB) is composed of specialized brain microvascular endothelial cells (BMECs) located in brain capillaries^1^ and has been the focus of intense study both *in vivo* and *in vitro* due to the large number of neurodegenerative diseases that are associated with BBB impairment. As a primary transport site between the blood and the brain, the BBB is a crucial regulator of brain homeostasis, maintaining both functional and biochemical connections with surrounding vascular support cells (pericytes, glia) and responding with vasoconstrictive or dilative signaling according to neuronal metabolic need.^2^ Disruption to the normal function of the BBB leads to a number of downstream and potentially irreversible consequences for overall brain health and cognition.^3^

While biochemical and cell-cell contributions to BBB function and changes in disease have been extensively explored, mechanical contributions to BBB regulation are less examined. Numerous studies have shown that among older adults, arterial stiffness, primarily measured by elevated aortic pulse wave velocity (PWV), is associated with worse cerebral small vessel disease, such as increased white matter hyperintensities^4^ and worse cognitive outcomes.^5^ Arterial stiffening increases vascular tone, which reduces the capacity for dynamic autoregulation and contributes to reductions in regional cerebral blood flow, which has been shown in older individuals with elevated PWV.^6^ Further, adverse cognitive outcomes associated with arterial stiffening are usually linked to the hemodynamic consequences of impaired cerebral blood flow and increased pulsatile stress reaching the cerebral microvessels due to reduced aortic compliance. Thickening of the basement membrane also occurs in smaller cerebral arterioles in older adults with and without dementia, suggesting a co-occurrence of matrix remodeling and altered hemodynamics in smaller brain vessels.^7^ While changes to the capillary basement membrane are not possible to resolve with current neuroimaging technology, advancing age and Alzheimer’s disease (AD) are associated with reduced microvascular density and increased presence of “string vessels,” which are capillaries that have lost their endothelium and cannot transmit blood.^7,8^ Overall, there is a long-standing consensus that advanced age and vascular risk factors drive structural alterations to the small cerebral arterioles and capillaries which either precede or co-occur with AD and related dementias. However, tracing the explicit cellular mechanisms that connect BBB dysfunction to these mechanical changes remains an area of active investigation.

Owing to difficulties controlling and measuring shear stress and substrate stiffness *in vivo*, efforts to assess the role of mechanical forces on endothelial function have primarily focused on *in vitro* models. Multiple *in vitro* studies of peripheral and brain endothelial cells have demonstrated sensitivity to fluid shear stress magnitude and flow profiles (e.g. oscillatory vs pulsatile flow).^9–11^ Application of physiological shear stress improves markers of BMEC identity *in vitro*^12^ and decreases barrier permeability compared to static cultures.^11^ By comparison, excessively high shear stress deteriorates markers of barrier function^9^ and promotes endothelial dysfunction in human umbilical vein endothelial cells.^10^ While observations of peripheral endothelial cells *in vivo* show consistent elongation and alignment in the direction of blood flow,^13^ *in vitro* BMEC models have demonstrated inconsistent response to FSS. A recent 2D system applying rotational shear stress to multiple BMEC lines (stem-cell derived, immortalized, primary) showed that BMECs align perpendicular to the direction of flow, while other BMEC models show a lack of flow alignment and reduced cell turnover in BMECs compared to peripheral endothelial cells.^14–16^ Together, these conflicting data collected with different *in vitro* systems suggest the need for further investigation. To our knowledge, all *in vitro* BBB models that incorporate fluid flow utilize either extremely soft, compliant substrates^15^ or non-compliant surfaces coated in a thin layer of ECM.^11,17,18^ A study utilizing peripheral endothelial cells showed that cells grown in static culture on tissue culture plates, in comparison to a soft 4 kPa hydrogel, began to transition to a mesenchymal state, behaving similarly to cells treated with a known chemical trigger of endothelial-to-mesenchymal transition (EndoMT).^19,20^ While demonstrating clear endothelial responsivity to extremely stiff versus soft substrates, tissue culture plates lack any mechanical similarity to physiological tissue.^21^ Work from our lab with induced pluripotent stem cell-derived BMEC (iBMEC)-like cells demonstrated a reduction in intracellular actin stress fibers on intermediate stiffness polyacrylamide (PA) hydrogels (20 kPa), although there was no difference in junctional protein width or tortuosity.^22^ By contrast, a recent study utilizing two different stem cell lines for derivation of iBMEC-like cells demonstrated an increase in the continuous junction pattern of ZO-1 staining on 15 and 194 kPa PA hydrogels, compared to 1 kPa and 2.5 kPa PA hydrogels.^23^ These studies demonstrate a clear sensitivity of iBMEC-like cells to substrate stiffness in regard to junctional and cytoskeletal protein distribution, but further work is needed to reach a consensus on the magnitude and impact of these changes on BBB dysfunction. Additionally, no studies have sought to clarify the collective influence of substrate stiffness and shear stress on BMEC phenotypes. Here, we show that primary human BMECs adopt an inflammatory signature when cultured on 30 kPa gelatin hydrogels relative to 6 kPa gelatin hydrogels, and that this inflammatory signature could be partially attenuated by exposing the BMECs to modest shear stress. These results provide new insight into mechanical regulation of BMEC inflammation involving integrated responses to both the substrate and external flow.

## 3. Methods

### 3.1 Preparation of GEL hydrogel and crosslinking solution

Gelatin (GEL) solutions (recorded by w/v%) were prepared by dissolving powdered porcine gelatin (300g bloom Type A, Sigma, G1890) in ultrapure water and heating to 40°C. After hydration, GEL was filter-sterilized using a heated 0.22 μm Steriflip filter (Millipore) and maintained at 40°C for same day experiments. GEL was kept at 4°C for up to 1 week after sterile preparation and reheated as necessary. Microbial transglutaminase (mTG) solution (recorded by w/v%) was prepared by dissolving powdered mTG (TG, Meat Glue, MooGloo) in ultrapure water, which was heated to 40°C and sterile filtered. mTG solution was kept at 40°C until used. A 10% (w/v) mTG solution was added directly to 6.5% GEL at a (v/v) ratio of 3:100, gently mixed, and quickly dispensed into the desired location. For the 15% GEL, a 20% (w/v) mTG solution was added directly to the GEL at a (v/v) ratio of 3:100, gently mixed, and quickly dispensed into the desired location. The 6.5% GEL was allowed to crosslink for 24 hours in the cell culture incubator. For the 15% GEL samples, additional crosslinking solution (20% mTG) was added to the surface of the GEL for an additional 24 hours before being washed 3x with PBS. GEL samples used for cell culture were further coated with an extracellular matrix (ECM) solution before use, as described later.

### 3.2 Rheological determination of shear modulus (G*) on crosslinked GEL

Rheological data were collected on crosslinked hydrogel samples to determine the complex shear modulus (G*) when samples were exposed to low stress. Complex shear modulus was calculated from the storage modulus (G’) and loss modulus (G’’) in the linear viscoelastic region, as shown in Equation 1. Hydrogel discs were prepared by crosslinking samples in 3D printed molds (2 mm deep, 20 mm diameter) designed to fit the diameter of the 20 mm rheometer plate. After 24 hours, samples were removed from the molds with a metal spatula, washed 3x in PBS to remove residual crosslinker, blotted with a kimwipe, and placed on the bottom plate of a parallel plate rheometer (AR 2000, TA Instruments). After lowering the plate head to make complete contact with the top surface of the sample (no slip confirmed by a lack of free rotation of the scan head), an oscillation amplitude strain sweep experiment was run (once per sample) to determine the shear modulus. Sample temperature was maintained at 37°C, and conditions were as follows: strain range of 0.00645% to 6.45%, angular frequency of 1 rad/sec, and torque of 10 to 100 µN x m. After data collection was complete (1 cycle per sample), complex shear modulus (G*) was calculated. Following the methods outlined in a previous publication characterizing the mechanical properties of gelatin,^24^ the complex Young’s Modulus (E*) was calculated using a Poisson’s ratio (υ) of 0.451 (assuming there is negligible difference between our hydrogel υ and native gelatin at similar w/v%) as seen in Equation 2. Average E* was then calculated over the linear viscoelastic region, identified by the strain range over which G* remained relatively constant, according to Equation 2. Please note that GEL substrates are referenced by their Young’s modulus from these rheological measurements (6 kPa or 30 kPa) for the remainder of the manuscript.

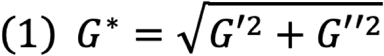

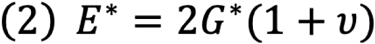

### 3.3 Dynamic mechanical analysis for determination of Young’s Modulus (E) on crosslinked GEL

Dynamic mechanical analysis (DMA) was used to directly measure the Young’s Modulus (E) of hydrogel samples. A Q series 800 DMA (TA Instruments) was used to analyze crosslinked GEL samples. GEL samples were prepared by crosslinking in a small petri dish and punching out circular samples with a 6 mm biopsy punch. Circular samples were kept hydrated under sterile conditions until measurements were completed. All measurements were completed at 37°C under a compressive strain ramp of 0.5% strain/min. Young’s Modulus was determined from the slope of the linear region of the stress vs strain curve from the strain ramp data.

### 3.4 Scanning electron microscopy analysis of crosslinked GEL samples

For scanning electron microscopy, lyophilized hydrogels were mounted on double-sided carbon tape and attached to a Ted Pella pin mount. The top surfaces of the gels were leveled using a scalpel. After gold sputtering, the samples were observed using a scanning electron microscope (Zeiss Merlin) at an accelerating voltage of 5 kV and magnification of 300X. Images were processed in ImageJ to quantify pore size.

### 3.5 Rheological determination of media viscosity

Rheological measurements of the dynamic viscosity of cell culture media were performed on an AR 2000 rheometer (TA Instruments) using a 40 mm parallel plate attachment and 20 µm measuring gap. Before each measurement session, the instrument was mapped with the 40 mm plate and the zero point was re-calibrated. To measure dynamic viscosity, a steady state flow experiment was completed with a shear rate ramp from 0.1 to 100 Hz at 37°C following 1-2 minutes of sample conditioning at the experiment temperature. The rheological measurements of the average dynamic viscosity across a shear rate of 0.1 to 100 Hz for endothelial cell media, with or without 3% (w/v) 70 kDa dextran (Sigma, C31390), were measured. Equation 3 describes the relationship between shear stress (, fluid viscosity (**µ**), and the shear rate () under Newtonian conditions.

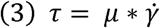

### 3.6 Measurement of volumetric flow rate

Volumetric flow rate was measured at the inlet of devices (directly after a fluid splitter) using two different liquid flow sensors (SLI-0430 & SLI-2000, Sensirion) for low (< 5AU) and high (> 5AU) flow rates, with sampling rate of approximately 16.7 Hz. Continuous measurements of volumetric flow were recorded for a minimum of 60 seconds at each splitter outlet (a tubing segment of equal length was attached to the outlet not being measured). Average flow rate was calculated across triplicate measurements from 2 full periods of pulsatile flow for each pump speed measured. Minimum and maximum flow rates were averaged across each sample measurement for calculations of the full flow rate range.

### 3.7 Primary brain microvascular endothelial cell culture

Primary human BMECs were obtained from Cell Systems (ACBRI 376) and maintained in Endothelial Cell Growth Media (C-22110, Sigma) up to passage 9. ACBRI 376 cells are isolated from healthy donor tissue and confirmed positive for CD31 and VWF, per the supplier information. For all 2D experiments, BMECs were subcultured onto the ECM-coated GEL at a density of 1×10^6^ cells/mL and allowed to adhere for 24 hours before induction of flow. Endothelial cell media was supplemented with 3% (w/v) 70 kDa dextran to increase media viscosity. All experiments were completed using ACBRI 376 cells from the same vial. No media changes were performed following the start of each 48-hour experiment.

### 3.8 Preparation and operation of ibidi devices

Each ibidi µ-slide I luer 3D (“ibidi”) device is composed of three separate hydrogel wells in series that are sealed by a glass coverslip adhered to the top surface of the slide (87176, Ibidi). Each device is perfused by a single inlet port connected via silicone tubing (MFLX96410-13, VWR) to a peristaltic pump system (Ismatec). After all hydrogel components were prepared, 16 µL of GEL + mTG solution was pipetted into each well and a rectangular glass coverslip was adhered to the top of the device. Ibidi devices were moved to cell incubators and allowed to crosslink for 24 hours in a humidified petri dish. For the 30 kPa GELs, enhanced crosslinking was completed by hydrating the surface of the wells with additional 20% mTG (compared to water for the 6 kPa GEL). Following crosslinking, an ECM solution was introduced to the apical surface of the hydrogel wells via the inlet ports. ECM was freshly made before each use and was composed of 50 ug/mL human collagen IV (Sigma, C5533), 25 ug/mL bovine fibronectin (Sigma F1141), and 1x Penicillin-Streptomycin (P/S) in DMEM/F12. After approximately 24 hours of ECM incubation, BMECs at 80-90% confluency were collected with TripLE and resuspended in appropriate media supplemented with 10 µM Rho kinase inhibitor (Y-27632, Tocris) and 1x P/S and introduced to the inlet port of the device. Cells were allowed to adhere to the GEL surface for 24 hours before connecting devices to the perfusion system and initiating fluid flow. For all perfusion experiments, the endothelial cell media was supplemented with 3% (w/v) 70 kDa dextran which has been previously shown to promote microvessel stability *in vitro.*^25^ To reduce the possibility of bubble formation, sterile media was acclimated to the incubator conditions for 4 hours before being connected to the perfusion system and a bubble trap (Darwin Microfluidics, LVF-KBT-L-A) was inserted directly upstream of the ibidi compartment. Ibidi devices were perfused with an 8-channel peristaltic pump (Ismatec) that remained outside the incubator, with tubing fed through a custom-printed rubber plug in the back of the cell culture incubator. Adhered cells were allowed to acclimate to low flow conditions (10-43 uL/min) for 24 hours before pump speed was increased to the final flow rate (e.g. 248 uL/min) for 48 hours.

### 3.9 Preparation of static GEL cultures

Static conditions were completed by culturing BMECs on GEL substrates cast directly into wells of a 24-well glass bottom plate. GEL (6.5% or 15%) was prepared as previously described and approximately 300 µL of GEL + mTG was carefully pipetted against the bottom of the well. Immediately after dispensing, plates were manually rotated to ensure even coating of the GEL and any bubbles were aspirated before the GEL thermally crosslinked. Empty wells were filled with water to prevent the GELs from drying out, and the plates were moved to cell incubators and allowed to crosslink for 24 hours. For the 30 kPa GELs, enhanced crosslinking was completed by hydrating the surface of the wells with additional 20% mTG (compared to water for the 6 kPa GEL). Following crosslinking, GELs were incubated with the extracellular matrix (ECM) solution described in the previous section. After approximately 24 hours of ECM incubation, BMECs were cultured on GEL substrates following previously described protocol. Cells were allowed to adhere to the GEL surface for 24 hours. Following 24 hours of cell adhesion in endothelial media supplemented with ROCK inhibitor, media was changed to endothelial cell media supplemented with 3% (w/v) 70 kDa dextran (identical to perfusion media) and cells were kept in the incubator for an additional 48 hours without further media changes.

### 3.10 Immunocytochemistry

Each sample was fixed and stained following standard protocols. Cell culture media was removed, and cells were washed 2x with PBS. Cells were then fixed with 4% PFA for 10 minutes and washed 3x with PBS (5 minutes per wash). Cells were then washed 2x with PBS-T (0.1% Triton-X) and allowed to incubate for 10 minutes. Following permeabilization, cells were incubated with blocking buffer (5% goat serum, Invitrogen #500627) for 10 minutes followed by incubation with primary antibody for either 3 hours at room temperature or overnight at 4°C. After primary antibody incubation, cells were washed 3x with PBS (10 minutes per wash) and incubated with the appropriate combination of secondary antibodies and conjugated antibodies or other fluorescent markers. Secondary/conjugated antibodies were incubated at room temperature for 3 hours in the dark or overnight at 4°C on a plate rocker. After a final PBS wash (3x, 10-minute incubations), cells were stained with DAPI for 30-60 minutes, washed 3x with PBS, and immediately imaged. Representative images showing select BMEC identity markers (PECAM-1, Claudin-5) were minimally preprocessed in FIJI to improve visualization of cell borders. DAPI, F-Actin, and Claudin-5 channels were auto enhanced and subjected to background subtraction. All cell samples were imaged on a Leica LAS X epifluorescent microscope (10x lens) or a Zeiss LSM 710 Confocal Microscope (10x lens). Secondary antibody only controls were utilized. See **Supplementary Table 1** for a list of all antibodies used in these experiments.

### 3.11 Analysis of Cell Morphology

Cell morphology analyses were completed using Cell Profiler (version 4.3.8) by analyzing F-actin labeled cells in representative images from each condition that were captured on a Leica LAS X epifluorescent microscope.^26^ For each condition, a minimum of 3 biological replicates (3 separate wells or 3 ibidi devices) with 3 technical replicates (ROI selected from non-overlapping regions in the same well) were analyzed. Image preprocessing was completed using FIJI (version 2.14.0) wherein all images were cropped to the same area (712 x 532 µm), split into individual channels, and each channel was converted to an 8-bit greyscale image and saved as a .tiff. For FSS conditions, greyscale images were contrast enhanced using the auto “Enhance Contrast” function on FIJI with 0.35% pixel saturation to improve resolution of cell borders. All prepared images were loaded into the Cell Profiler pipeline and the Identify Primary Objects module was used to identify the cell nucleus from the DAPI (blue) channel by minimum cross-entropy thresholding. Following the nuclear identification, the cell body was identified with the Identify Secondary Objects module. All objects touching the image border were discarded. For both modules, the threshold correction factor and smoothing scale were manually adjusted and tested against multiple images in each set before applying to the whole image set. Quantification of cellular and nuclear shape was completed by averaging morphological features from the MeasureObject module by the total cell count from the sum of all technical replicates. Cell Profiler pipelines are included in the Supplemental Information.

### 3.12 ICAM-1 surface levels

Quantification of ICAM-1 levels was completed using FIJI (version 2.14.0) for image preprocessing and Cell Profiler (version 4.3.8) for quantification of F-actin and ICAM-1 from images captured on a LSM 710 confocal microscope. Microscope settings were kept constant across imaging sessions. Representative images of each condition underwent preprocessing to improve visualization of cellular structures. Each representative image was cropped to the same ROI area (425.1 x 425.1 µm) split into individual channels in FIJI and the auto adjust brightness/contrast feature was applied to the F-Actin and DAPI channels for improved visualization. For image quantification of ICAM-1 positive regions, raw images (n=3 biological replicates, each with 3 technical replicates taken at different regions in the same well) were cropped to the same ROI (425.1 x 425.1 µm) and channels were split and the LUT was changed to greyscale. To aid in identification of cell bodies, the “Enhance Contrast” function was applied to the red (F-Actin) channel only, with pixel saturation at 0.35% and the “Normalize” option selected. All cropped images were then saved as 16-bit .tiff files and imported into Cell Profiler. All preprocessed images were loaded into the Cell Profiler pipeline and the Identify Primary Objects module was used to identify the cell nucleus from the DAPI channel by minimum cross-entropy thresholding. Following the nuclear identification, the cell body was identified with the Identify Secondary Objects module and minimum cross-entropy thresholding. ICAM-1 positive regions were identified with the Identify Primary Objects module, using Otsu thresholding. All objects touching the image border were discarded. For both modules, the threshold correction factor and smoothing scale were manually adjusted and tested against multiple images in each set before applying to the whole image set. The total area (in pixels) occupied by cells in each representative image was measured by the MeasureObject module. The total ICAM-1 positive area per image was similarly measured and used to calculate the % ICAM-1 positive area over the total area covered by cells for each image. The reported % coverage for each biological replicate is the average of all three representative images. Cell Profiler pipelines are included in the Supplemental Information.

### 3.13 Cellular distribution of ZO-1

Quantification of ZO-1 junctional versus cytoplasmic + nuclear intensity for static samples was completed using FIJI (version 2.14.0) using methods described previously.^14^ Representative images from each condition were opened and cropped to the same dimensions (425 x 425 µm). The line drawing tool was set to 4-pixel width and used to trace a straight section along the cell border and the average intensity was recorded. The circle drawing tool was used to select for the cell cytoplasm + nucleus, with care taken to avoid potential overlap with cell junctions. Three representative cells were selected for measurement per image. A total of 9 images were analyzed, comprised of 3 separate wells and 3 technical replicates (images taken in separate locations) per well. All cells chosen for measurement did not share a border and all parts of the cell membrane had to be easily identifiable without any adjustment of pixel intensity. Reported data represent the average of all measurements across all technical replicates per well. Fluorescence intensity is quantified by pixel value from raw images.

### 3.14 Bulk RNA sequencing

RNA was extracted from ibidi devices or static plates with TRIzol incubation for 10 minutes, followed by purification with a Direct-zol RNA miniprep kit (Zymo, #R2050). Concentration of purified RNA (resuspended in 20-25 µL of RNAse-free water) was quantified with a NanoDrop and stored at -80°C until needed for sequencing. RNA samples were submitted to the Vanderbilt Technologies for Advanced Genomics (VANTAGE) core for sequencing and a LabChip GX was used to ensure RNA was of sufficient quality before proceeding with sequencing. cDNA libraries were prepared with a stranded mRNA (polyA-selected) library preparation kit. Next generation sequencing was performed at paired-end 150 bp on an Illumina NovaSeq X Plus with a targeted 50M reads per sample. Demultiplexed FASTQ files were aligned using quasi-mapping transcriptome alignment with *Salmon*.^27^ A decoy-aware whole genome transcriptome index for alignment was created with the GENCODE GRCh28 primary assembly genome and v46 transcripts (GRCh38.primary_assembly.genome.fa, gencode.v46.transcripts.fa). Following quantification of paired-end reads with *Salmon*, count data was analyzed following the Bioconductor pipeline in R (version 4.4.2). Transcriptomic count data was imported and converted to gene-level quantification with *tximeta*^28^ and differential gene expression was quantified with the *DESeq2* and results were compared for select conditions using a grouping variable (e.g. by FSS or hydrogel).^29^ Additional packages used: *glmpca*, *paletteer*, *tidyr, apeglm,*^30^ *magrittr, dplyr, tidyverse*. For data visualization of all samples, the dataset was pre-filtered to remove genes with 0 or very low counts (< 10). DEGs were computed from paired comparisons between sample conditions comparing either the hydrogel substrate or flow rate: (1) 6 kPa vs 30 kPa GEL at 1.7 dyne FSS, (2) 6 kPa vs 30 kPa GEL at 0 dyne FSS, (3) 0 dyne vs 1.7 dyne FSS for 6 kPa GEL, and (4) 0 dyne vs 1.7 dyne FSS for 30 kPa GEL (see Supplemental Information for single condition versus all other conditions). Heatmap plots showing the normalized count data or log2foldchange values between paired results using *pheatmap* package. Volcano plots were created with *ggplot2* where significance cut off values are p-adjusted (padj) < 0.01 and log2foldchange > 1 for upregulated genes and < 1 for downregulated genes.

### 3.15 Pathway analysis and enrichment map visualization

Pathway analysis was completed using DEG datasets generated with *DESeq2* following the protocol described previously.^31^ Gene set enrichment analysis (GSEA) (version 4.3.0)^32^ from preranked lists was completed with the most recent version of the Gene Ontology, Reactome, and KEGG Medicus gene sets from the MSigDB database (c5.go.bp.v2024.1.Hs.symbols.gmt, c2.cp.kegg_medicus.v2024.1.Hs.symbols.gmt, c2.cp.reactome.v2024.1.Hs.symbols.gmt) for 1000 permutations in combination with the provided Chip platform for Ensembl gene ID conversion (Human_Ensembl_Gene_ID_MSigDB.v2024.1.Hs.chip). To improve understanding and visualization of significant pathways, enrichment networks were created from GSEA results using Cytoscape (version 3.10.2) with the EnrichmentMap, AutoAnnotate, and WordCloud apps.^31^ For all enrichment maps, cut-off values of p < 0.005, FDR q-value < 0.01, and overlap of 0.5 were used to generate the map, and cluster labels were individually examined and modified from the WordCloud label based on an examination of the gene sets, highlighting the common themes.

### 3.16 Analysis of conditioned media and cell lysate samples

Conditioned media was collected from each sample and centrifuged, and the supernatant was stored at - 20°C until further analysis. ELISA was completed for soluble ICAM-1 (sICAM-1, Thermo Fisher, BMS241, extra sensitive) following manufacturer’s instructions. For conditioned media samples from the static conditions, each individual well is considered as a single biological replicate. For the +FSS conditions, each conditioned media sample is pulled from the media that was perfusing two ibidi devices, due to the flow loop set up that contained a fluid splitter. Following the collection of conditioned media, cells were washed 1x with PBS and incubated with 100 uL of protease/phosphatase inhibitor cocktail diluted 1:50 in RIPA buffer for 30 minutes on ice. Following buffer incubation, samples were washed 3-4x with the RIPA cocktail, transferred to clean microcentrifuge tubes, and centrifuged at 4°C for 15 minutes at 12000 rpm. Supernatant was carefully collected and moved to clean microcentrifuge tubes and stored at -80°C until needed. Total protein per sample (for conditioned media and cell lysate) was determined using a Pierce BCA protein assay kit and microplate reader (Cytation 3, BioTek). Triplicate samples for each condition were submitted to Eve Technologies Corporation for multiplexed analysis of a human cytokine panel, HDF15, using Luminex xMAP technology. All Luminex samples were diluted to the same concentration of total protein (119 µg/mL) to account for differences in total cell surface area and measured in duplicate.

### 3.17 Statistical Analysis

All statistical analysis was performed in GraphPad Prism (version 10.4.0) or RStudio. Figure legends include the statistical tests utilized for each experiment.

## 4. Results

### 4.1 Choice of hydrogel substrates

An overview of the experimental set up describing hydrogel conditions, BMEC subculture, and induction of fluid shear stress (FSS) is shown in **Figure 1A**. To develop this system to simultaneously study responses to shear stress and substrate stiffness, we first needed to select a hydrogel system with mechanical properties similar to organ tissues.^21^ Gelatin (GEL) was chosen as the hydrogel substrate for all experiments due to the relative ease of handling, low cost, tunable stiffness, and excellent cell adhesion properties. It is also composed of similar amino acid composition as collagen, the primary component of the vascular basement membrane. By tuning the gelatin concentration and crosslinker density, we developed two GEL hydrogels with a bulk Young’s modulus of 6.4 ± 1.6 kPa (6 kPa GEL) and 29.6 ± 5.3 kPa (30 kPa GEL), respectively (**Figure 1B**). Our primary measure of GEL stiffness was determined by rheological measurements of GEL samples (**Figure 1C** shows example of strain sweep), which directly measures the shear modulus of elasticity (G) by the storage and loss moduli. The shear modulus can be directly related to the Young’s modulus by the use of Poisson’s ratio, following previous characterization of pure gelatin hydrogels.^24^ We also directly measured the Young’s modulus of our two final GELs by the use of dynamic mechanical analysis (DMA) under compressive stress (**Figure 1D**). While DMA was less precise than rheological measurements, they are in line with our rheological data, showing an order of magnitude difference in the Young’s modulus (4.69 kPa vs 55.6 kPa) as shown in **Table 1**. To determine if changing the gelatin concentrations and crosslinker density resulted in substantially different pore sizes, we imaged representative hydrogel samples by scanning electron microscopy but did not measure a significant difference in pore size between samples (**Supplementary Figure 1**). Due to the fluid shear forces that are the focus of this study, we believe that G* is the more relevant elastic modulus to describe the GEL behavior and deformability under 2D flow. However, most *in vitro* models that include substrate stiffness report Young’s modulus. In the rest of this manuscript, we refer to the GEL substrates as 6 kPa vs 30 kPa.

**Figure 1:**
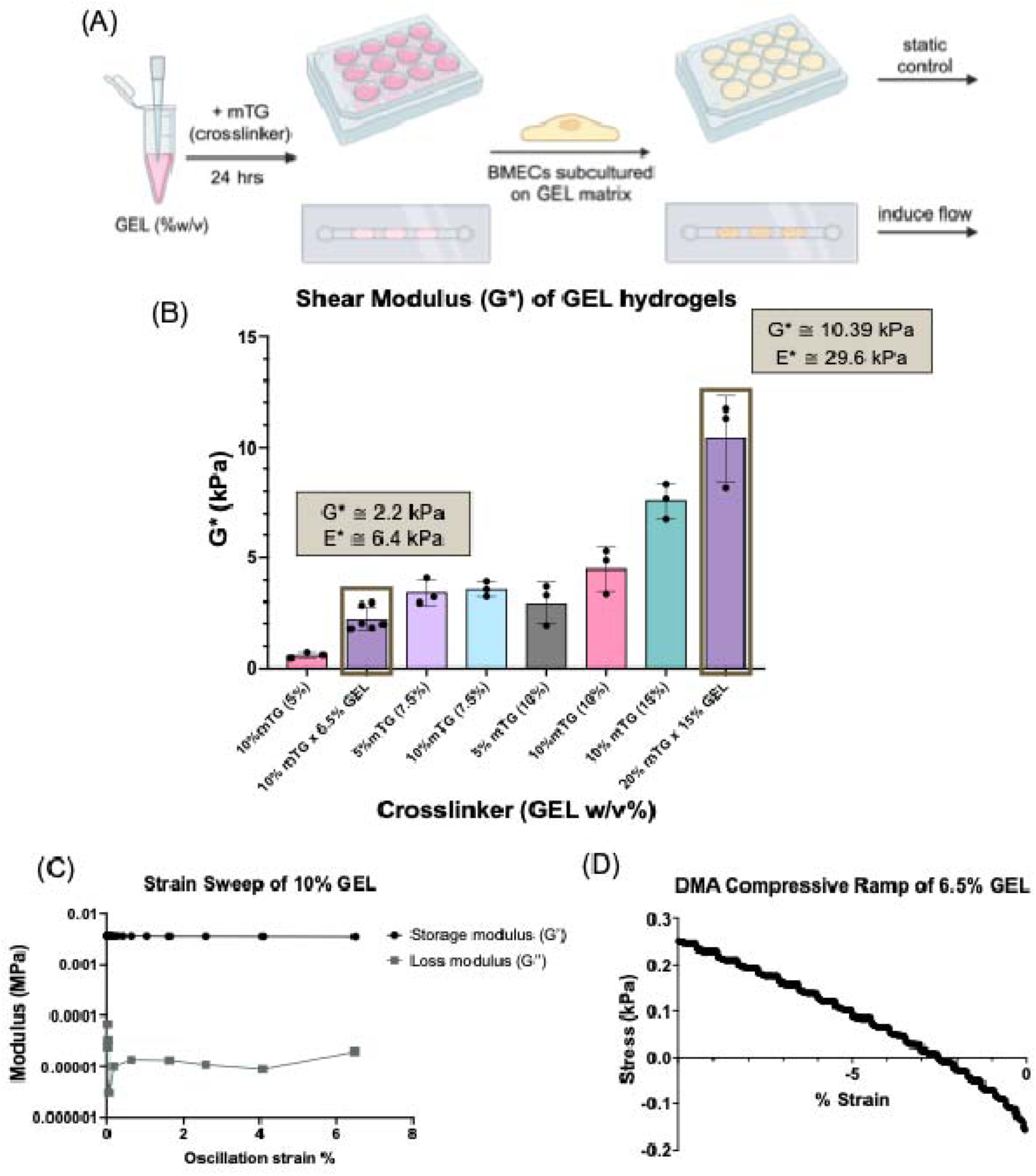
Experimental set up and characterization of hydrogel substrate stiffness. (A) General experimental set up for *in vitro* models with GEL substrate. (B) Influence of GEL (w/v) and crosslinker concentration on shear modulus, as measured by rotational rheology. A minimum of n=3 independent replicates were measured per GEL condition (mean ± SD). (C) Example of strain sweep measurement of 10% GEL sample on rheometer, showing storage and loss modulus. (D) Example of DMA compressive ramp for direct measurement of Young’s modulus from the slope of the linear region. For each GEL sample, n=3 independent replicates were measured and mean ± SD is reported in Table 1. DMA measurement data are found in Supplementary Figure 2.

**Table 1:** Determination of Shear Modulus and Young’s Modulus of GEL hydrogels.

|  |  |  |  |
| --- | --- | --- | --- |
| Measurement type | Sample | Young's Modulus, $E^*$ (kPa) | Shear Modulus, $G^*$ (kPa) |
| DMA (compressive stress) | 6.5% GEL, 0.5% strain/min | $4.69 \pm 1.47$ | N/A |
| | 15% GEL, 0.5% strain/min | $56.55 \pm 22.83$ | N/A |
| Rheometer (shear stress) | 6.5% GEL, $\nu = 0.5$ | $6.4 \pm 1.57$ | $2.2 \pm 0.55$ |
| | 15% GEL, $\nu = 0.5$ | $29.6 \pm 5.29$ | $10.39 \pm 1.94$ |

### 4.2 Choice of shear stress conditions

FSS is a consequence of the tangential frictional force of blood as it moves parallel to the vascular endothelium. Due to the cardiac cycle, blood flow is inherently pulsatile, and we therefore chose to use a peristaltic pump system to achieve regular pulsatile flow for our experiments (**Figure 2A-B**). Assuming Newtonian behavior, shear stress (***τ***) is a product of the fluid viscosity (**µ**) and the shear rate 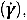 which is determined from the vessel geometry and volumetric flow rate. Newtonian fluids are defined by a linear relationship between applied force and deformation, where viscosity is constant for a material at a given temperature and pressure. In addition to the presence of serum in the cell culture media, we added a low concentration of dextran when perfusing media through devices, which has been shown to help stabilize 3D microvessels *in vitro* and increases the viscosity of the media.^25^ While previous studies have shown that the addition of serum to cell culture media results in the transition to non-Newtonian behavior,^33^ the majority of *in vitro* systems make the simplification to Newtonian physics for the purpose of shear stress calculations. We therefore chose to use an average viscosity across 1-100 Hz (**Figure 2C**) to calculate the average FSS experienced by cells in the ibidi system. For this study, we chose to focus on the interaction of substrate stiffness with the absence or presence of low, pulsatile flow with an average FSS of 1.7 dyne/cm^2^ which is within the range of estimates for physiological FSS *in vivo*.^34^ FSS calculations are found in **Table 2**.

**Figure 2:**
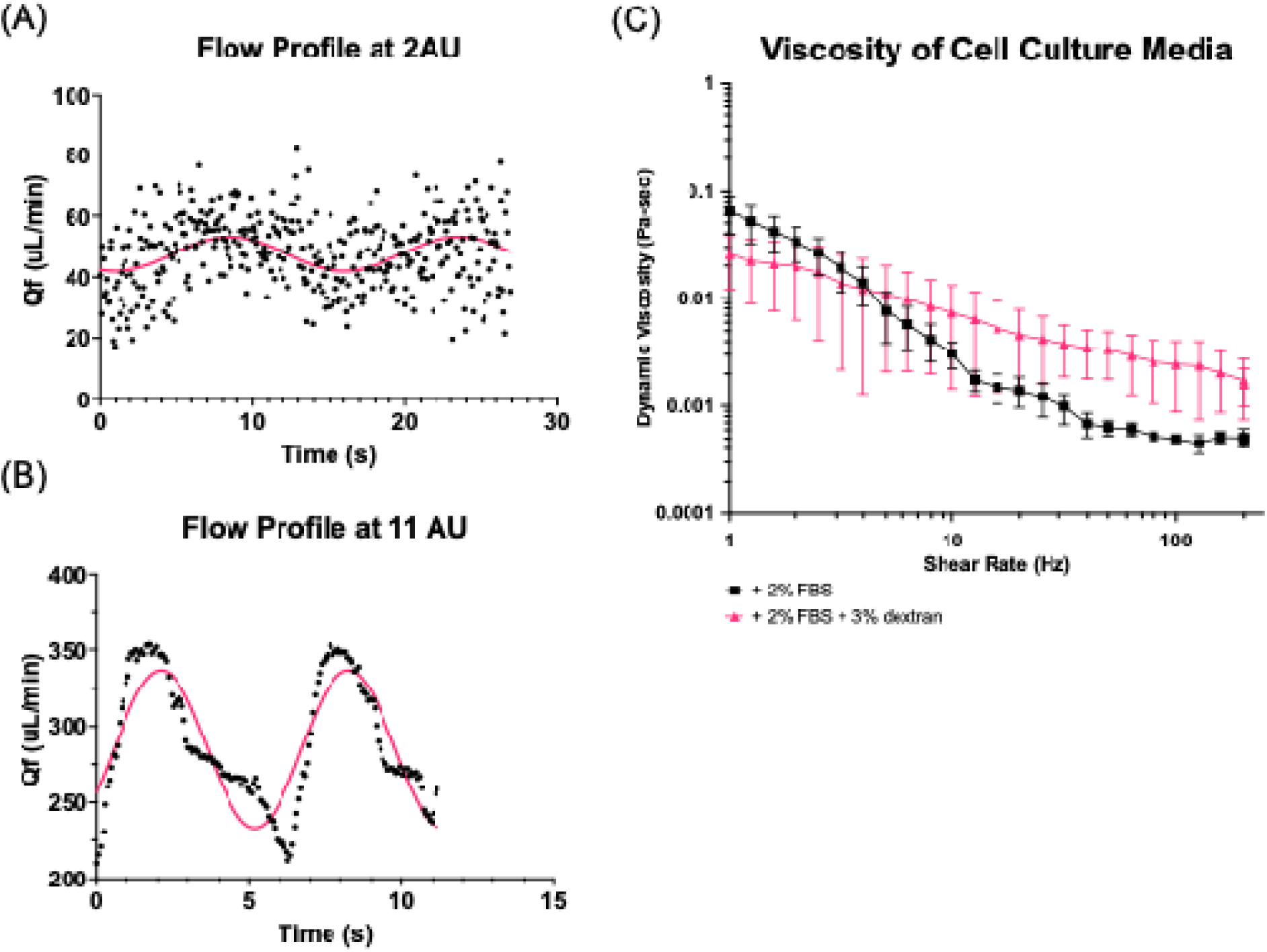
Pulsatile flow profile and measurement of media viscosity for FSS calculations. (A-B) Volumetric flow profile sampled at 2 AU (pump speed, arbitrary units) and 11 AU that was measured directly at the inlet to the ibidi device. Pink line represents a best fit sine wave to aid in visualization of the pulsatile nature of the fluid flow. A single representative measurement is shown for each condition, and flow rates for FSS calculations are the average of n=3 independent measurements. (C) Viscosity of endothelial cell media supplemented with 2% serum and with or without 3% (w/v) of 70 kDa dextran. Each media sample was measured in triplicate and data are presented as mean ± SD. While both media compositions exhibit a tendency towards non-Newtonian behavior (viscosity being dependent on shear rate), for the sake of simplicity we have assumed the average viscosity of the media to be a constant from 1-100 Hz and calculated the FSS according to Equation 3. See Table 2 for flow rates (average of n=3) and corresponding FSS within the ibidi device.

**Table 2:**
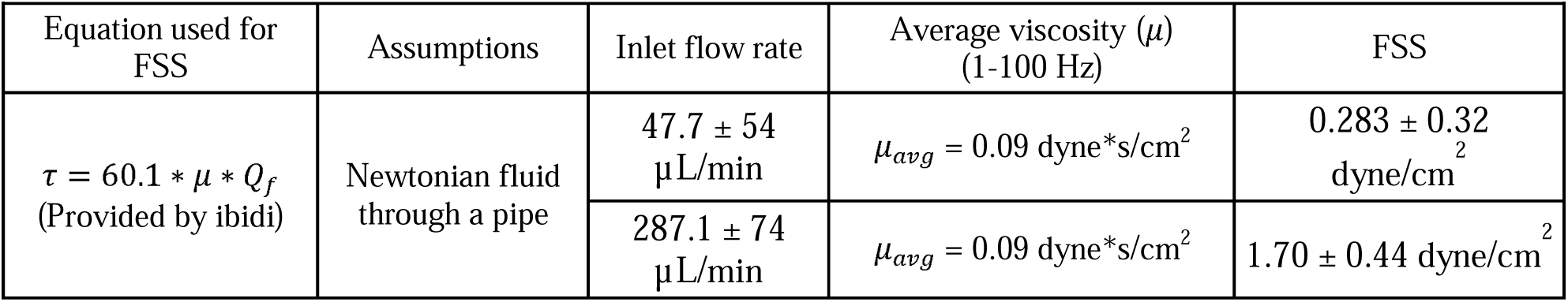
Calculation of FSS in ibidi devices.

| **Table 2: Calculation of FSS in ibidi devices** |  |  |  |  |
| --- | --- | --- | --- | --- |
| Equation used for FSS | Assumptions | Inlet flow rate | Average viscosity ($\mu$) (1-100 Hz) | FSS |
| $\tau = 60.1 * \mu * Q_f$<br>(Provided by ibidi) | Newtonian fluid through a pipe | 47.7 ± 54 μL/min | $\mu_{avg} = 0.09$ dyne*s/cm<sup>2</sup> | 0.283 ± 0.32 dyne/cm<sup>2</sup> |
| | | 287.1 ± 74 μL/min | $\mu_{avg} = 0.09$ dyne*s/cm<sup>2</sup> | 1.70 ± 0.44 dyne/cm<sup>2</sup> |

### 4.3 BMEC morphology and junctional markers are sensitive to mechanical forces

We first sought to characterize general BMEC responses to substrate stiffness with or without application of FSS. Primary BMECs were confirmed to express CD31 and Claudin-5 after subculture on both 6 kPa and 30 kPa GEL substrates (**Figure 3A-B**). Under static conditions, we observed a significant decrease (p=0.03) in the junctional/cytoplasmic+ nuclear localization of ZO-1—a crucial intercellular scaffolding protein which connects tight junction complexes to the actin cytoskeleton and is associated with BBB integrity^35^—for BMECs cultured on the 30 kPa GEL compared to the 6 kPa GEL (**Figure 3C-D**). We also observed small differences to cellular morphology as a consequence of both substrate stiffness and FSS (**Figure 3E-K**). Cell size on 30 kPa GEL and 6 kPa GEL was unchanged by static conditions (p=0.2445) but significantly higher under 1.7 dyne/cm^2^ FSS (p=0.003) on both substrates (**Figure 3F**). Total cell count (by number of nuclei) also decreased significantly under 1.7 dyne/cm^2^ FSS (p=0.0282) for both the 6 kPa and 30 kPa substrates (**Figure 3G**). Addition of FSS significantly decreased cellular circularity (Form Factor; p<0.0001) and increased both Major Axis (p=0.0088) and Minor Axis (p=0.0003) length for both substrates (**Figure 3H, J, K**). Cellular orientation, where 45° indicates random orientation, was increased in the direction of flow across both substrates under FSS (p=0.0013), indicating that substrate identity contributed to the alignment response under FSS (**Figure 3I**). Overall, these results indicate BMECs exhibit collective sensitivity to mechanical forces from FSS and the underlying substrate, motivating deeper exploration into cell states.

**Figure 3:**
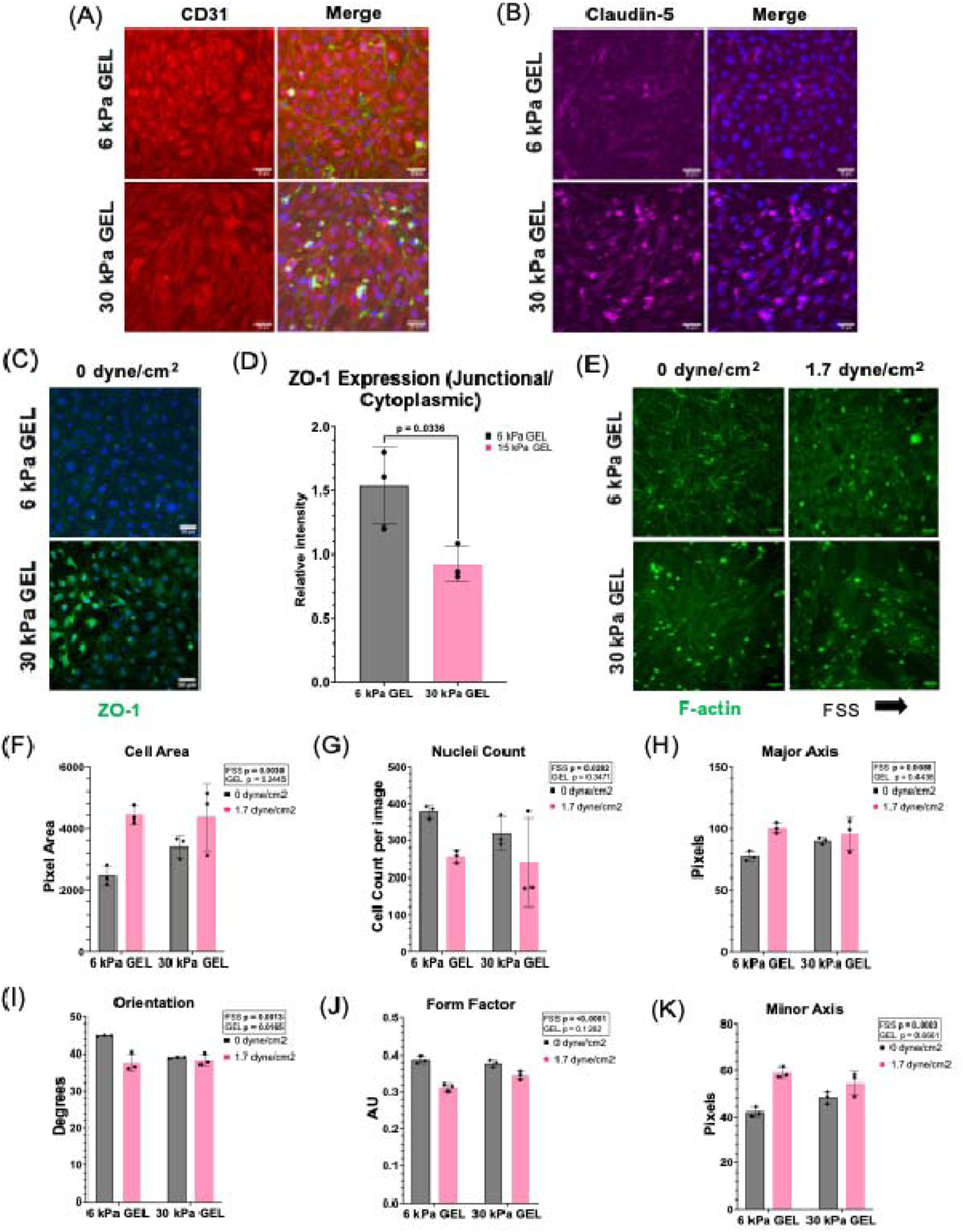
Influence of substrate stiffness and FSS on junctional protein localization and cellular morphology. (A) BMECs subcultured on 6 kPa and 30 kPa GEL substrates under static conditions express CD31 (red), merged image includes F-actin (green) and DAPI (blue). (B) BMECs subcultured on 6 kPa and 30 kPa GEL substrates under static conditions express claudin-5 (magenta). Merged images include DAPI (blue). (C) Representative images showing ZO-1 (green) and DAPI (blue) on 6 kPa and 30 kPa GELs under static conditions. (D) Quantification of ZO-1 intensity ratio between cell junctions and cell cytoplasm (including nucleus). Data are presented as mean ± SD from 3 biological replicates. Each replicate is calculated from measurements on nine total images across 3 technical replicates. Significance was determined by an unpaired student’s t-test. (E) Representative images of F-actin stain used to identify cell shape for cell morphology analyses. (F-K) Quantification of cell morphology characteristics using Cell Profiler. Each datapoint represents a single biological replicate that is the average from all technical replicates divided by the total cell count in the ROI. Data are presented as mean ± SD. Significance and interaction terms by GEL and FSS were determined by two-way ANOVA. Representative images were minimally preprocessed in FIJI to improve visualization of cell borders and reduce background. Scale bars are 50 *µ*m for all images.

### 4.4 Transcriptomic data and pathway analysis

To better understand the combined impact of substrate stiffness and FSS on cellular function, we performed bulk RNAseq (**Figure 4A**) on mRNA collected from BMECs exposed to the following conditions: (1) 6 kPa GEL at 0 dyne/cm^2^ FSS, (2) 30 kPa GEL at 0 dyne/cm^2^, (3) 6 kPa GEL at 1.7 dyne/cm^2^, and (4) 30 kPa GEL at 1.7 dyne/cm^2^ FSS. Following processing of FASTQ files (**Figure 4A**) for each sample, we performed principal component analysis (PCA) (**Figure 4B**) and found similar clustering for all samples from the same condition. For differential expression analysis, we used a grouping variable to assign each condition with a FSS and substrate identifier and directly compared gene expression in a pairwise manner. After filtering the data to remove genes with counts < 10, we had 17032 total genes for differential expression analysis. Paired comparison between the 6 kPa and 30 kPa GELs under 0 dyne/cm^2^ FSS had the smallest number of total DEGs (2572 genes) out of all paired sample comparisons (> 7000 genes), which aligns well with the variance displayed in the PCA plot (**Figure 4C**). A snapshot of the top 10 enriched pathways from gene set enrichment analysis (GSEA) shows the most significant cellular pathways that are upregulated in the 6 kPa GEL compared to the 30 kPa GEL are related to cell cycle checkpoints, chromosome alignment, and nuclear division (1.7 dyne/cm^2^ FSS) and protein translation (0 dyne/cm^2^ FSS) (**Figure 4D**). By comparison, the majority of upregulated pathways in the 30 kPa GEL compared to the 6 kPa GEL are related to chemokine receptor binding, IL-10 signaling, and immune cell migration across both FSS states. Under static conditions, BMECs on the 6 kPa GEL had upregulated transcriptomic profiles related to immune response and protein translation pathways when compared to BMECs exposed to 1.7 dyne/cm^2^ FSS (**Figure 4F**). By comparison, BMECs on the 6 kPa GEL exposed to 1.7 dyne/cm^2^ FSS had upregulated pathways primarily related to cell-cell and cell-ECM adhesion. Finally, for BMECs cultured on the 30 kPa GEL, the static condition demonstrated upregulated pathways related to both cell cycle checkpoints and immune-related signaling compared to the 1.7 dyne/cm^2^ FSS condition, with predominant negative regulation for multiple pathways related to TGF-*β* signaling (**Figure 4G**).

**Figure 4:**
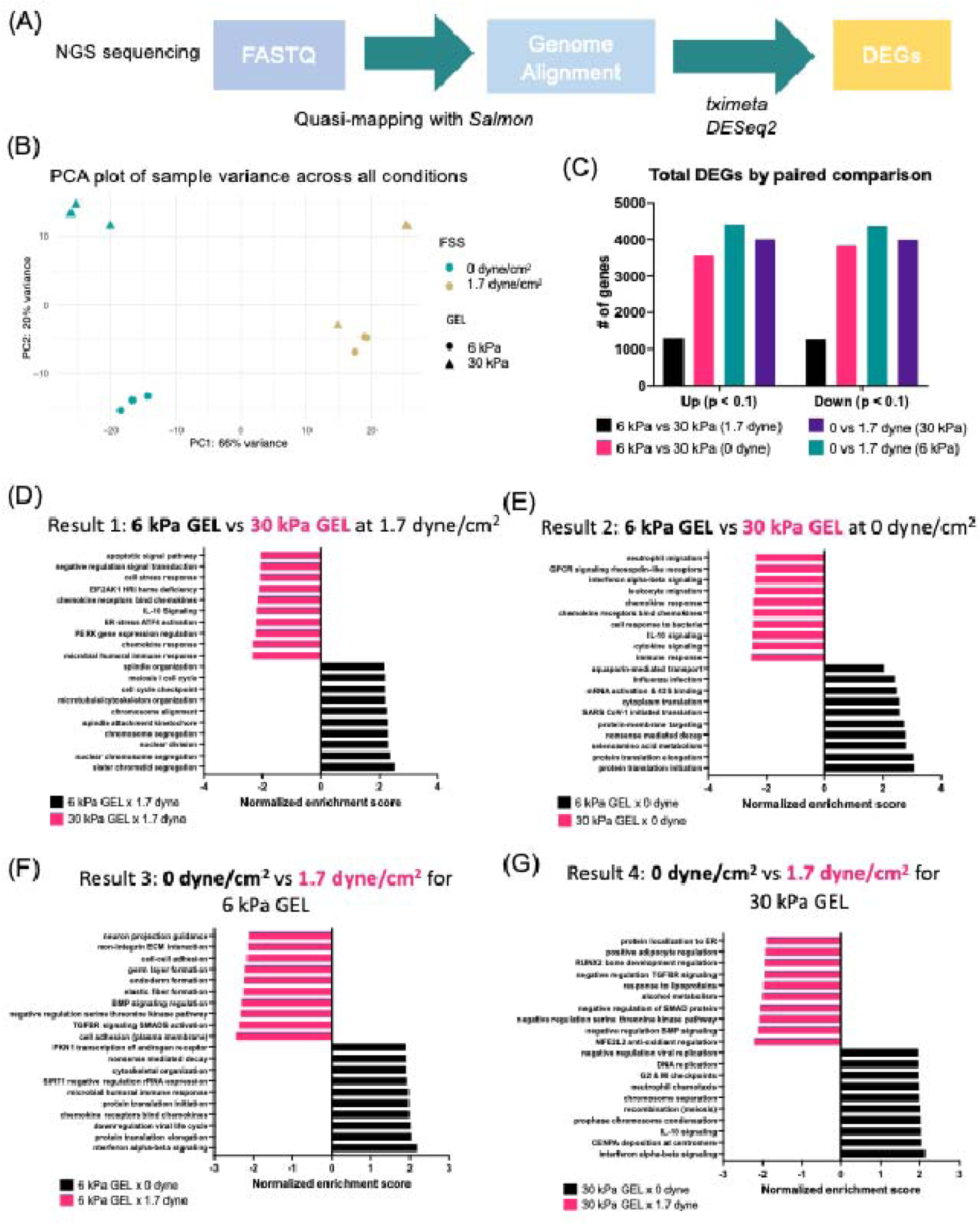
Overview of bulk RNAseq analyses across GEL substrate and FSS. (A) Simplified overview of pipeline used to process raw RNAseq data for determination of DEGs. (B) PCA plot displaying variance between all samples (n=3 biological replicates per condition) with expected clustering by sample condition. Samples are identified by shape for GEL substrate (circle = 6 kPa GEL, triangle = 30 kPa GEL) and color for FSS (turquoise = 0 dyne/cm^2^, brown = 1.7 dyne/cm^2^). Note that two of the samples for the 1.7 dyne/cm^2^ x 30 kPa GEL condition (brown triangles) showed extremely similar transcriptomic signatures and are clustered on top of each other. (C) DEGs were determined by pairwise comparison of conditions with identical substrate or FSS, resulting in 4 sets of results. Total DEGs for all comparisons (p < 0.1) are shown by upregulated genes (UP in the first term) and downregulated genes (UP in the second term). (D-G) Top 10 most significant pathways from GSEA ranked by normalized enrichment score (NES) for each set of paired results. Because we utilized 3 gene set databases for our pathway analysis, highly overlapping/duplicate pathways were manually removed from these data. See Supplementary Figure 3 for data on DEGs and Supplementary Figure 5 for volcano plots of top 10 pathways comparing transcriptome from each condition with all other samples.

While pathway analysis indicated several cellular pathways related to cell cycle regulation, cell adhesion, and immune activation were affected by substrate and/or FSS conditions, we chose to focus on the immune activation response, which showed notable differences for BMECs cultured on the 30 kPa GEL compared to the 6 kPa GEL. We first examined the transcriptomic response under static conditions, where the upregulation in inflammatory-related pathways was the most prominent on the 30 kPa GEL compared to the 6 kPa GEL (**Figure 5A**). From the pathway analysis results, we selected a mid-sized gene set (GO:1990868) related to the cellular response to chemokines and used this gene set to selectively annotate volcano plots for comparisons (**Figure 5B-E**). These annotations confirm that many of the significant differentially regulated genes for the 30 kPa GEL were indeed related to the cellular response to chemokines (**Figure 5B-C**). When comparing directly across FSS states for the 30 kPa GEL condition, the majority of chemokine receptor genes were upregulated in the static condition, suggesting a role of FSS dampening the inflammatory response (**Figure 5E**). Overall, cellular pathways related to inflammatory response were most upregulated in the 30 kPa GEL at 0 dyne/cm^2^ FSS, and specific genes related to the cellular response to chemokines showed significant upregulation on the 30 kPa GEL at 0 dyne/cm^2^ FSS when compared to 6 kPa GEL (**Figure 5B**) or compared to 1.7 dyne/cm^2^ FSS (**Figure 5E**). **Table 3** shows the specific adjusted p-values (padj) and log2FoldChange of the annotated gene set.

**Figure 5:**
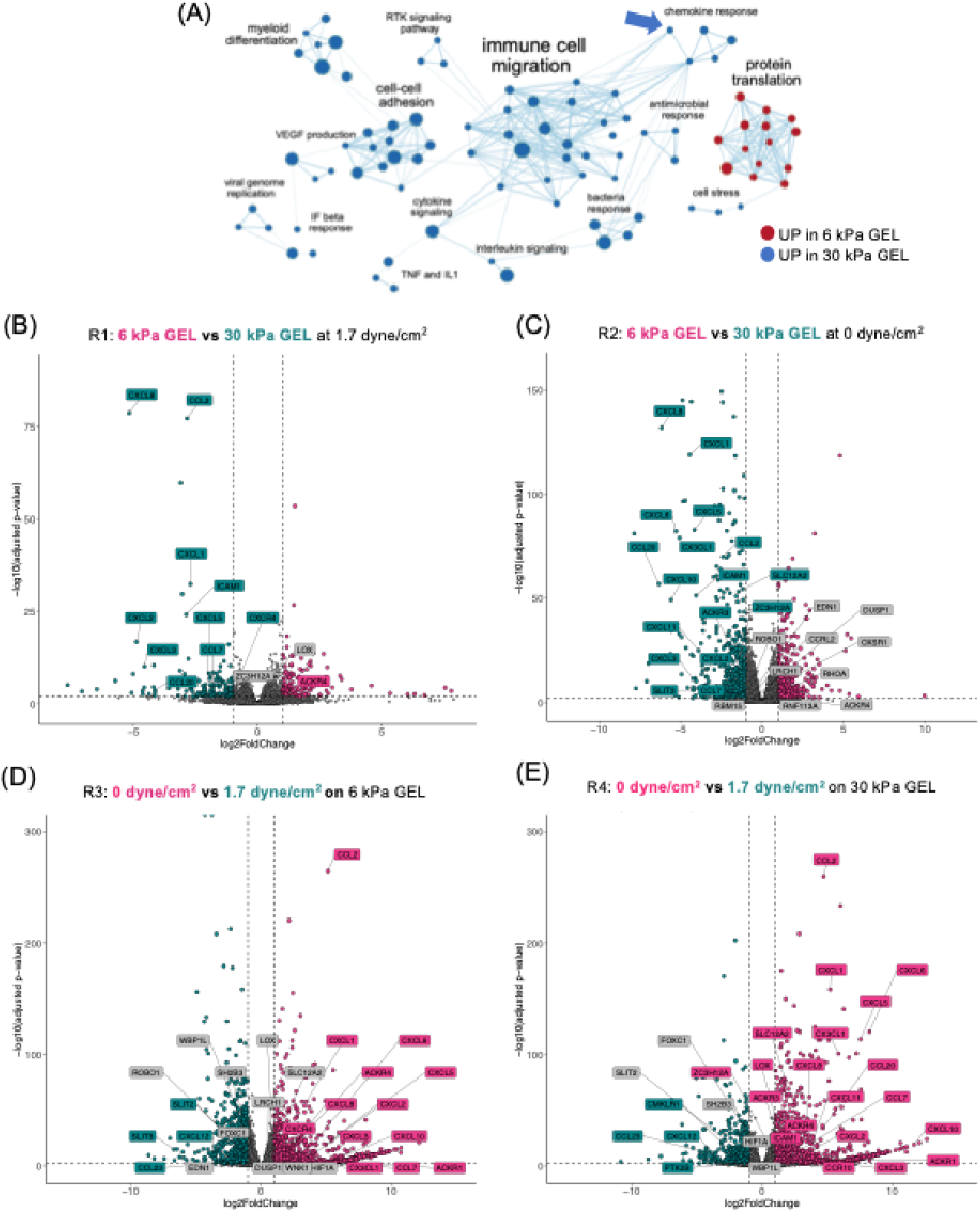
Pathway analysis reveals inflammatory signature in BMECs cultured on 30 kPa GELs compared to 6 kPa GELs. (A) Simplified enrichment map of significantly upregulated cellular pathways BMECs on 30 kPa GELs (blue) versus 6 kPa GELs (red) under static conditions. Node significance threshold was set by NES at FDR q-value < 0.005 and edge cutoff (similarity) was set to 0.5. Singleton nodes and clusters that were not connected to any other clusters were manually removed from enrichment map. Blue arrow indicates GS used for volcano plot annotations (response to chemokine, GO:1990868). (B-D) Volcano plots of paired comparisons between experimental conditions. Significance cutoffs (dashed lines) represent adjusted p-value (padj) < 0.01 and log2foldchange ± 1. Response to chemokine (GO) was used as the reference gene set, with the addition of *ICAM1*, to selectively annotate the volcano plot. Gene symbol labels are colored pink, indicating significant upregulation in the first term of the comparison, or turquoise, indicating upregulation in the second term of the comparison. Genes colored in grey have padj < 0.1 but log2foldchange between -1 and 1. See Table 3 for the og2foldchange and p-adj values for all annotated genes.

**Table 3:** Summary statistics of annotated genes in volcano plots (GO:1990868)

|  |  |  |  |  |  |  |  |  |
| --- | --- | --- | --- | --- | --- | --- | --- | --- |
| Gene ID | L2FC(R1) | padj(R1) | L2FC(R2) | padj(R2) | L2FC(R3) | padj(R3) | L2FC(R4) | padj(R4) |
| ACKR1 | NA | NA | NA | NA | 3.97084177 | 0.00012469 | 4.27758048 | 2.79E-07 |
| ACKR3 | NA | NA | -2.0287624 | 1.32E-28 | NA | NA | 1.50028668 | 6.99E-16 |
| ACKR4 | 1.16230219 | 0.00192043 | 0.87599677 | 0.00621334 | 1.70479796 | 1.29E-08 | 1.99110338 | 1.03E-10 |
| CCL2 | -2.8471836 | 6.37E-78 | -2.4173296 | 1.06E-70 | 5.14049572 | 2.50E-265 | 4.71064164 | 1.47E-260 |
| CCL20 | -4.1370703 | 0.00023203 | -6.3374547 | 1.50E-57 | NA | NA | 4.65802097 | 1.27E-34 |
| CCL23 | NA | NA | NA | NA | -2.3952284 | 4.71E-05 | -4.4869246 | 6.39E-13 |
| CCL7 | -1.8205103 | 0.00787934 | -1.3325547 | 0.00028992 | 1.97769981 | 0.00024057 | 1.48974427 | 0.00032816 |
| CCR10 | NA | NA | NA | NA | NA | NA | 1.74678591 | 0.00043571 |
| CCRL2 | NA | NA | -0.4276568 | 3.89E-06 | NA | NA | NA | NA |
| CMKLR1 | NA | NA | NA | NA | NA | NA | -1.9033979 | 1.02E-07 |
| CX3CL1 | NA | NA | -5.0733617 | 1.03E-79 | 1.17924489 | 7.51E-05 | 5.85776948 | 2.14E-101 |
| CXCL1 | -2.7070704 | 4.62E-33 | -4.4477545 | 1.32E-119 | 3.47181547 | 2.21E-58 | 5.2124995 | 5.49E-159 |
| CXCL10 | NA | NA | -5.5896452 | 5.70E-50 | 5.33185068 | 7.37E-05 | 11.6526326 | 7.27E-20 |
| CXCL11 | NA | NA | -3.9017086 | 1.18E-25 | NA | NA | 3.88163657 | 4.47E-24 |
| CXCL12 | NA | NA | NA | NA | -3.5258343 | 1.28E-10 | -3.9906289 | 4.16E-13 |
| CXCL2 | -4.8759751 | 2.00E-17 | -3.6994448 | 3.76E-14 | 4.98986788 | 5.30E-20 | 3.81333754 | 5.73E-15 |
| CXCL3 | -4.5579297 | 7.99E-11 | -4.2636803 | 5.24E-12 | 3.42725901 | 2.74E-07 | 3.13300954 | 7.22E-07 |
| CXCL5 | -2.0593518 | 0.00700333 | -4.1632436 | 1.72E-83 | 5.42534252 | 1.85E-23 | 7.52923435 | 2.99E-114 |
| CXCL6 | NA | NA | -5.2883839 | 6.83E-83 | 4.26698629 | 1.11E-24 | 8.23269569 | 4.43E-121 |
| CXCL8 | -5.1699841 | 2.10E-79 | -6.1762567 | 1.20E-132 | 2.4822504 | 4.80E-19 | 3.48852301 | 6.02E-43 |
| CXCR4 | -1.0091704 | 0.00185519 | NA | NA | 1.86015599 | 2.75E-12 | NA | NA |
| DUSP1 | NA | NA | -0.5038147 | 0.00707156 | -0.4911753 | 0.00736563 | NA | NA |
| EDN1 | NA | NA | -0.7658404 | 0.00121839 | -0.9014615 | 8.20E-05 | NA | NA |
| FOXC1 | NA | NA | NA | NA | -0.5666166 | 9.65E-10 | -0.9985584 | 3.17E-28 |
| HIF1A | NA | NA | NA | NA | 0.50343197 | 0.00054679 | 0.74313745 | 1.94E-07 |
| ICAM1 | -2.8810025 | 7.68E-25 | -4.0598437 | 4.63E-52 | NA | NA | 1.60407227 | 1.01E-08 |
| LOX | 0.3526617 | 0.00018146 | NA | NA | 0.85262598 | 8.31E-28 | 1.06017465 | 3.32E-42 |
| LRCH1 | NA | NA | 0.44492278 | 5.40E-09 | 0.54181915 | 1.02E-12 | NA | NA |
| OXSR1 | NA | NA | -0.2843452 | 0.00082228 | NA | NA | NA | NA |
| PTK2B | NA | NA | NA | NA | NA | NA | -1.285634 | 0.00148264 |
| RBM15 | NA | NA | -0.4330178 | 0.00592008 | NA | NA | NA | NA |
| RHOA | NA | NA | 0.26593088 | 0.0082223 | NA | NA | NA | NA |
| RNF113A | NA | NA | 0.40654789 | 0.00945908 | NA | NA | NA | NA |
| ROBO1 | NA | NA | -0.767154 | 4.05E-09 | -0.540144 | 4.96E-05 | NA | NA |
| SH2B3 | NA | NA | NA | NA | -0.6312986 | 2.39E-20 | -0.4426169 | 2.83E-10 |
| SLC12A2 | NA | NA | -1.4583082 | 6.41E-54 | 0.53930653 | 5.04E-08 | 1.75413072 | 2.13E-77 |
| SLIT2 | NA | NA | NA | NA | -1.0464699 | 8.70E-09 | -0.670019 | 0.00044996 |
| SLIT3 | NA | NA | -4.2445118 | 1.62E-06 | -6.6768951 | 1.72E-15 | NA | NA |
| WBP1L | NA | NA | NA | NA | -0.5178708 | 2.08E-13 | -0.2257546 | 0.0032011 |
| WNK1 | NA | NA | NA | NA | -0.4720953 | 0.00615164 | NA | NA |
| ZC3H12A | -0.7975861 | 0.0008463 | -1.6184899 | 1.87E-17 | NA | NA | 1.03992098 | 1.43E-07 |

Note: For results comparisons, R1 refers to 6 kPa vs 30 kPa GEL at 1.7 dyne/cm<sup>2</sup> FSS, R2 refers to 6 kPa vs 30 kPa GEL at 0 dyne/cm<sup>2</sup> FSS, R3 refers to 0 vs 1.7 dyne/cm<sup>2</sup> FSS for 6 kPa GEL, and R4 refers to 0 vs 1.7 dyne/cm<sup>2</sup> FSS for 30 kPa GEL. L2FC refers to log2foldchange.

### 4.5 Quantification of cytokine production in cell lysate

To build on the transcriptomic analyses, we collected cell lysate to test for the presence of elevated cytokine production. We included two additional conditions in these experiments: (1) BMECs cultured on a glass plate at 0 dyne/cm^2^ FSS and (2) 30 kPa GEL at 0.3 dyne/cm^2^ FSS. BMECs cultured on glass were used to compare cell responses on a non-compliant surface, and the reduced magnitude of FSS was used to determine if responses to FSS were binary or graded. A Luminex immunosorbent assay with a human cytokine panel (GM-CSF, IFN*γ*, IL-1*β*, IL-1RA, IL-2, IL-4, IL-5, IL-6, IL-8, IL-10, IL-12p40, IL-12p70, IL-13, MCP-1, TNF*α*) was used to quantify protein concentration in cell lysate samples from each condition (**Figure 6A-D**). These results demonstrated a significant increase in inflammatory cytokine production of IL-6, IL-8, MCP-1, and IL-1Ra in all 30 kPa GEL conditions. Specifically, IL-6 (p=0.0115), IL-8 (p=0.0033), MCP-1/CCL2 (p=0.0039), and IL-1Ra (p=0.0045) were significantly different across substrates (including glass samples). Interestingly, MCP-1 production appeared to be significantly affected by both substrate (p=0.0039), and FSS (p=0.0015) with very little production in all perfused conditions. Overall, BMECs cultured directly on glass showed very little intracellular cytokine production, suggesting an important role for the viscoelastic properties of the GEL substrate in inducing an inflammatory response. Concentration of IL-8 was highest in the static condition and significantly decreased in response to FSS (p < 0.0001 compared to 0.3 or 1.7 dyne/cm^2^ FSS) (**Figure 6E**). Similar patterns were observed in the 30 kPa static GEL compared to exposure to 0.3 or 1.7 dyne/cm^2^ FSS, which significantly reduced IL-1Ra (p=0.0049, p=0.0113) and MCP-1 (p=0.0018, p=0.0018) (**Figure 6F-G**). In contrast, there were no significant differences in IL-6 production between 30 kPa GEL conditions across FSS (**Figure 6H**). When comparing only between the 0.3 or 1.7 dyne/cm^2^ FSS conditions on the 30 kPa GEL (**Figures 6E-H**), there were no significant differences in cytokine production. Overall, cell lysate samples showed a similar trend to the transcriptomic data, with 30 kPa GEL samples showing elevated production of IL-8, IL-1Ra, MCP-1 (for static samples), and IL-6.

**Figure 6:**
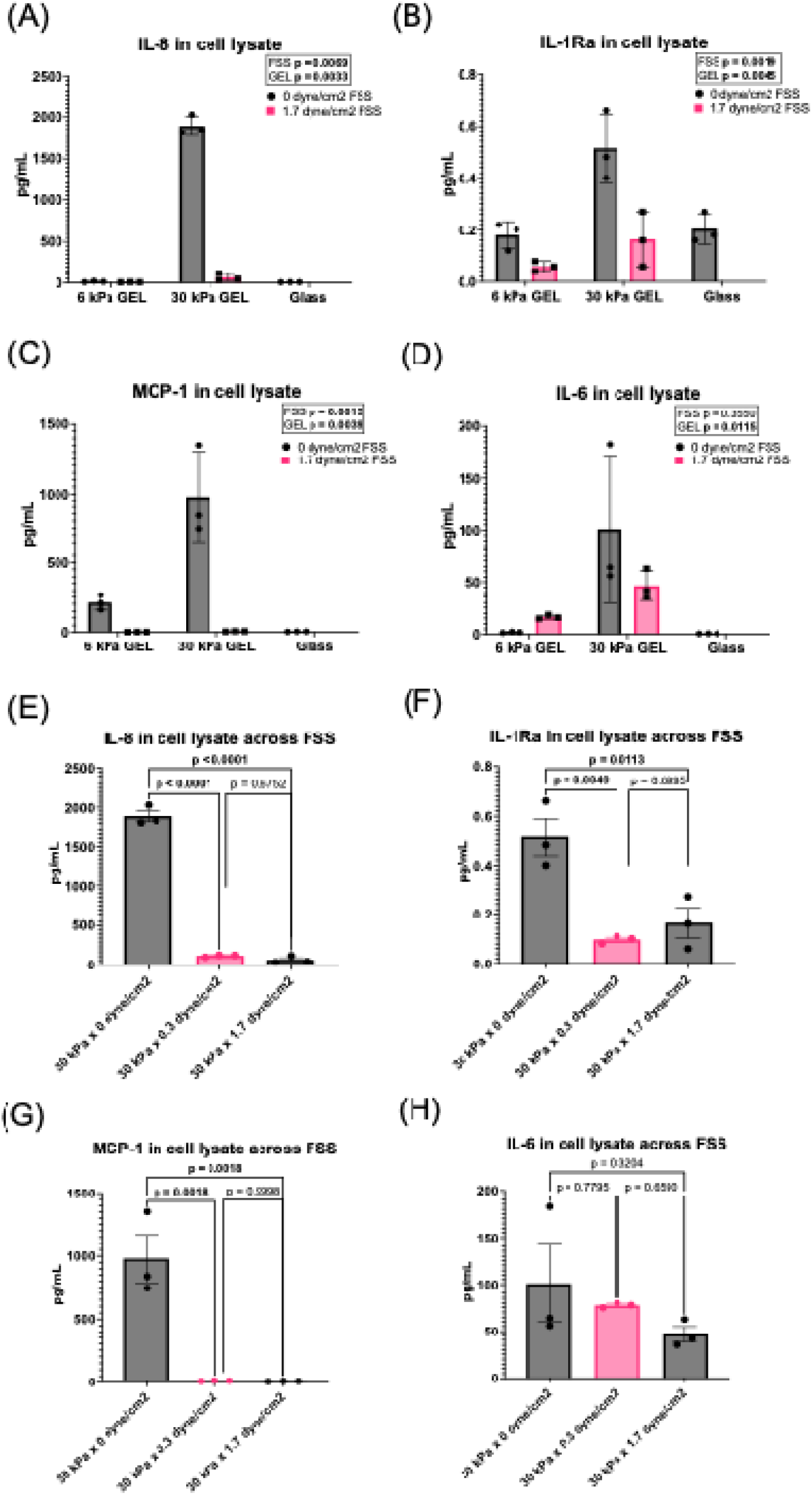
Cytokine levels in cell lysate. Select cytokine concentrations were measured in cell lysate using a luminex human cytokine panel (HDF15). All data are displayed as the average of duplicate measurements for n=3 biological replicates per condition (mean ± SEM). (A-D) Concentration of IL-8, IL-1Ra, MCP-1, and IL-6 grouped by GEL and FSS conditions. (E-H) Concentration of IL-8, IL-1Ra, MCP-1, and IL-6 at 0, 0.3, and 1.7 dyne/cm^2^ FSS for 30 kPa GEL substrate. Most of the cytokines in the panel were outside of the detection range (see Supplemental Information). Due to variations in GEL surface area and cell coverage between ± FSS experiments, all cell lysate samples were diluted to the same total protein count before analysis. Significance was determined via two-way ANOVA for grouped comparisons (panels A-D) and by one-way ANOVA for comparisons across 30 kPa GELs (panels E-H).

### 4.6 Quantification of soluble ICAM-1 in conditioned media samples (ELISA)

Following our analysis of cell lysate samples by the human cytokine panel, we conducted targeted ELISA assays of conditioned media samples to look at the profile of soluble ICAM-1 (sICAM-1). Due to the large differences in media volume between experimental setups, we did not directly compare between different FSS states for these samples. We measured sICAM-1 in all conditioned media samples, as *ICAM1* was differentially regulated in the transcriptome datasets. Under static conditions, sICAM-1 levels were significantly higher for the 30 kPa GEL compared to the glass samples (p=0.0054) but not compared to the 6 kPa GEL (p=0.2818) (**Figure 7D**). For the 1.7 dyne/cm^2^ FSS conditions, sICAM-1 levels were near the kit detection limit (0.33 ng/mL) and thus were not included in final analysis (data not shown).

**Figure 7:**
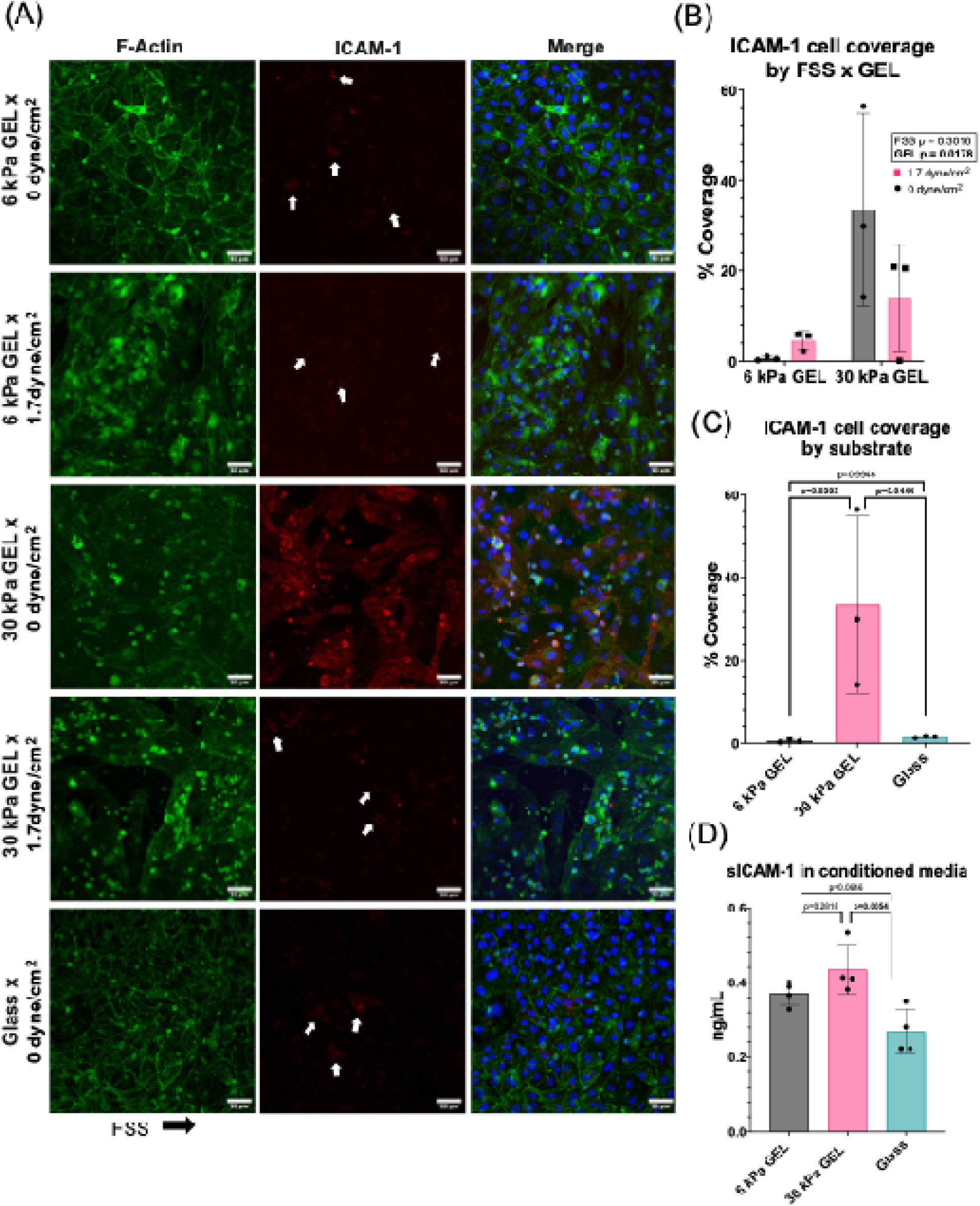
ICAM-1 levels on BMECs. (A) Representative images of F-actin (green) and ICAM-1 (red) under different FSS and substrate conditions. Arrows indicate visible ICAM-1 surface aggregation. (B) Percent coverage of positive ICAM-1 area over total cell coverage area for n=3 biological replicates per condition grouped by GEL and FSS (mean ± SD). (C) Percent coverage of positive ICAM-1 area for n=3 biological replicates across all substrates under static conditions (mean ± SD). (D) Concentration of sICAM-1 in conditioned media samples from 0 dyne/cm^2^ conditions (n=3 biological replicates, mean ± SD). BMECs cultured directly on glass were included for comparison in panels C-D. Significance determined by two-way ANOVA (panel B) and one-way ANOVA (panels C-D).

### 4.7 ICAM-1 surface expression across different GEL substrates and upon exposure to FSS

Based on our transcriptomic data showing an upregulation of *ICAM1* in BMECs cultured on the 30 kPa GEL, as well as increased levels of sICAM-1 for BMECs cultured on 30 kPa GEL versus glass, we quantified the surface coverage of ICAM-1 across all conditions. BMECs on 6 kPa GEL displayed few ICAM-1 positive regions regardless of exposure to FSS (**Figure 7A**). In contrast, BMECs cultured on the 30 kPa GEL at 0 dyne/cm^2^ FSS displayed significantly higher surface levels of ICAM-1, a trend which continues (though less dramatically) for the samples subjected to 1.7 dyne/cm^2^ FSS (p=0.0178) (**Figure 7B**). We also compared ICAM-1 surface levels across different substrates (including glass) under 0 dyne/cm^2^ FSS (**Figure 7C**) and found that the 30 kPa GEL had significantly higher ICAM-1 expression than the 6 kPa GEL (p=0.0393) and glass (p=0.0444) substrates, despite greater variability between samples. In conclusion, surface expression of ICAM-1 was significantly higher in BMECs cultured on 30 kPa GEL across both FSS states compared to BMECs on a softer substrate (6 kPa GEL) and stiff, non-compliant substrate (glass).

## 5. Discussion

### 5.1 Modest increases in the stiffness of compliant substrates promote an inflammatory response in human BMECs

In this study, we were interested in determining how the stiffness of compliant, viscoelastic substrates would contribute to BMEC dysfunction. Cellular substrates *in vitro* are usually a combination of collagen I,^25^ fibronectin, or hyaluronic acid, which provide a closer analog than gelatin to the composition of the basement membrane *in vivo* and are similar to parenchymal stiffness^36,37^ but are limited in their capacity to achieve stiffnesses > 4 kPa.^24,25^ A recent study utilizing iBMEC-like cells found that increased collagen I crosslinking (Young’s modulus of ∼4 kPa) improved barrier formation and stability in 2D cultures and 3D microvessels.^25^ Previous work in our lab demonstrated that increasing the stiffness of GEL hydrogels (0.3 kPa to 4.7 kPa) concurrently increased transendothelial electrical resistance, a surrogate for passive barrier strength.^22^ These studies motivated our interest in investigating the response of BMECs to wider range of stiffness that were informed by direct measurements of brain tissue and brain arterioles *ex vivo*.^39^ While we have previously used PA hydrogels^22^ to study BMEC responses to substrate stiffness, PA hydrogels lack higher-order structures and stress-strain dependent properties of native ECM that may influence cell behaviors.^21^

Our initial experiments in static cultures demonstrated a stiffness-mediated reduction of the junctional support protein, ZO-1, and translocation to the cytoplasm/nucleus (**Figure 3D**). This change suggested a potential disruption of BMEC barrier integrity, which would be in agreement with previous work.^22^ While we observed upregulated pathways related to cell-cell and cell-ECM adhesion in the 30 kPa GEL compared to the 6 kPa GEL, the main cellular response to the stiffer substrate was related to inflammatory pathways. Sustained vascular inflammation and endothelial activation is strongly associated with atherosclerotic plaque development,^40^ and elevated levels of inflammatory biomarkers in CSF are present early in Alzheimer’s disease and throughout disease progression.^41^ Our data show a consistent transcriptomic upregulation in inflammatory pathways related to chemokine and cytokine signaling and immune cell adhesion for BMECs cultured on 30 kPa GEL in both static culture and upon exposure to 1.7 dyne/cm^2^ FSS. Follow-up experiments from our transcriptomic analysis demonstrated elevated production of select inflammatory cytokines in BMECs cultured on the 30 kPa GEL, namely IL-6 and IL-8. Elevated levels of circulating IL-6 have been associated with worse cognitive performance and lower hippocampus and hypothalamus volumes in individuals with AD.^42^ IL-8 is the most potent chemoattractant of circulating neutrophils and is an important activator of the inflammatory response.^43^ It is expressed by multiple immune and vascular cell types, including endothelial cells, and is canonically upregulated in response to the reception of extracellular pro-inflammatory stimuli in the bloodstream.^43^ In the absence of additional cell types or introduction of pro-inflammatory cytokines in our experiments, our data suggest that *CXCL8* expression may be triggered by a cellular stress response to the stiffness of the underlying matrix. However, BMECs cultured directly on glass substrates did not show a similar upregulation in IL-8 expression, suggesting the importance of using compliant, viscoelastic substrates to capture more appropriate cell-matrix interactions.^21^ The upregulation in the production of pro-inflammatory chemokines and cytokines was accompanied by increased levels of surface ICAM-1, which would be expected to further promote the attachment and infiltration of circulating immune cells *in vivo*. This response is not without precedent, as peripheral endothelial cells show enhanced leukocyte adhesion after static culture on soft (0.2 kPa) or stiff (> 4 kPa) PA matrices, specifically identified by monocyte binding at ICAM-1 and IZ-actinin-4 clustering.^44^ Additionally, murine stem cells responded to increased hydrogel stiffness with elevated production of inflammatory cytokines.^45^ A murine triculture model studying effects of adipose tissue remodeling further showed that primary peripheral endothelial cells respond to stiffer collagen matrices (range from 0.292k–12.5 kPa) by upregulating *CCL2* and *IL6* mRNA expression.^46^ To our knowledge, our study is the first to tie BMEC inflammation to substrate mechanical properties.

### 5.2 Fluid flow modifies the effect of substrate stiffness on BMECs

Physiological FSS is an important mediator of vascular homeostasis. Multiple *in vitro* flow cell models have demonstrated that abnormal FSS patterns results in endothelial dysfunction for peripheral^10,47,48^ and brain endothelial cells.^14^ Our work suggests that the onset of physiological pulsatile flow dampens the observed inflammatory response in comparison to static conditions. Specifically, BMEC cultures subjected to 1.7 dyne/cm^2^ FSS on 30 kPa GELs had a reduced inflammatory signature when compared to 6 kPa GEL samples. To help identify the cause of this FSS-reduced response, we conducted a follow-up experiment where BMECs were subcultured on 30 kPa GEL and perfused at a lower 0.3 dyne/cm^2^ FSS. Analysis of cell lysate from the 0.3 dyne/cm^2^ FSS samples showed similar levels of IL-8, IL-1Ra, MCP-1, and IL-6 compared to the 1.7 dyne/cm^2^ FSS condition. These data may suggest that the presence or absence of FSS, rather than its magnitude, attenuates BMEC inflammatory responses on stiffer substrates. In contrast to the cytokines listed above, MCP-1 was significantly affected by both FSS and substrate stiffness. MCP-1 is a major chemoattractant, responsible for the recruitment of monocytes and macrophages *in vivo* and has been targeted for the treatment of various inflammatory-related diseases, such as atherosclerosis.^49^ These data suggest that production of MCP-1 may be stimulated by static cultures, potentially serving as an analog of stalled blood flow, which is associated with immune cell recruitment and remodeling. Previous work in an AD mouse model found that neutrophils were primarily responsible for stalling brain capillaries, and binding the neutrophil surface receptor Ly6G reduced the number of stalled capillaries and rescued cognitive impairment.^50^ The addition of FSS to our model provides evidence that the substrate stiffness mediated inflammatory response may be partially reduced in the presence of FSS.

## 6. Conclusion

In this study, we have shown that primary human BMECs generate an inflammatory response when cultured on compliant GEL substrates of moderate stiffness. By comparing cytokine production profiles across different cellular substrates, including BMECs cultured directly on glass, we also demonstrate that the fundamental nature of the substrate (i.e. viscoelastic hydrogel versus brittle solid) is a necessary consideration informing cellular response beyond simply the bulk stiffness.^21^ Further work will be necessary to help identify the mechanism(s) driving these responses. It is also important to consider substrate properties beyond stiffness measurements. While images of dehydrated hydrogels did not indicate significant differences in pore size, it is notoriously difficult to approximate pore sizes between dehydrated and hydrated samples. Further, increasing the gelatin concentration of our hydrogels likely increases the available integrin binding domains on the 30 kPa substrate, which may contribute to differences in the cell-ECM interaction between the 6 kPa and 30 kPa GELs. We are interested in conducting further studies on the dynamics of integrin-mediated cell-ECM attachment sites to understand how different cellular adhesion on each GEL substrate could be contributing to the observed inflammatory response. While the FSS values induced in this study were on the lower end for estimates of FSS *in vivo*,^13^ we were able to show that even small changes to FSS were sufficient to reduce the inflammatory response initiated by the 30 kPa substrate. We have designed our system to be easily modifiable to dynamic FSS environments, and future studies will build on the current hypothesis of stiffness-mediated inflammation by including the interaction of substrate stiffness with pathological FSS (> 40 dyne/cm^2^)^9^. Overall, our work provides evidence for a dynamic BMEC response to modest changes in their mechanical microenvironment, highlighting the role of vascular stiffening as a contributor to neurovascular inflammation.

## Abbreviations

FSS: fluid shear stress
BMEC: brain microvascular endothelial cells
GEL: gelatin hydrogel
PA: polyacrylamide hydrogel
PWV: pulse wave velocity
BBB: blood-brain barrier
ECM: extracellular matrix
DMA: dynamic mechanical analysis
AD: Alzheimer’s disease

## Ethics approval and consent to participate

Not applicable.

## Consent for publication

Not applicable.

## Availability of data and materials

RNA sequencing data are uploaded on ArrayExpress and can be accessed with accession number E-MTAB-15011.

## Competing Interests

Not applicable.

## Funding

This work was supported by funding from the Chan Zuckerberg Initiative (2019-191850, Ben Barres Early Career Acceleration Award to ESL), the National Institutes of Health (R01 NS110665 to ESL), and the National Science Foundation (601413, CAREER Award to ESL). AKY was supported by the Vanderbilt Interdisciplinary Training Program in Alzheimer’s Disease (T32 AG058524). AK was supported by a National Science Foundation Graduate Research Fellowship. DC was supported by the Integrated Training in Engineering and Diabetes (T32 DK101003). APL was supported by the Training Program in Environmental Toxicology (T32 ES007028). ALJ was supported by NIH grant K24 AG046373.

## Author Contributions

AKY and ESL were responsible for the initial conceptualization of this project. Formal analysis of data, methodology, data visualization, validation, and writing of the original manuscript were completed by AKY. Funding acquisition was provided by ESL. Supervision was provided by ESL and ALJ. Primary investigation was performed by AKY and HNM. AK performed SEM measurements, pore size quantification, and wrote the corresponding methods section. HK, HM, and AK provided guidance on methodology and data visualization for RNAseq and pathway analysis. DC provided guidance on conceptualization of experiments and study direction throughout the project. AL provided guidance on methodology for immunosorbent assays. AKY and ESL completed the majority of reviewing and editing the final manuscript. All authors were given the opportunity to review and approve the final manuscript before submission.

## Acknowledgements

Confocal microscopy was conducted in the Vanderbilt Cell Imaging Shared Resource core facility, which is supported in part by NIH grants P30 CA068485, P30 DK058404, and P30 EY008126. RNA sequencing was conducted in the Vanderbilt Technologies for Advanced Genomics core facility, which is supported in part by NIH grants 5UL1 RR024975, P30 CA068485, P30 EY008126, UL1 TR002243, and G20 RR030956. An award from the Vanderbilt Institute for Clinical and Translation Research (VICTR) to AKY (VR72110) was used to support some RNA sequencing costs—VICTR is supported by NIH grant UL TR002234. Scanning electron microscopy was performed in the Vanderbilt Institute for Nanoscale Science and Engineering. The authors are grateful to Dr. Leon Bellan and Romario Lobban for use of the Ares-G2 rheometer for GEL rheology measurements. We are also grateful to Dr. Scott Guelcher and Skyler Hornback for use of the AR2000 rheometer and helpful discussions regarding media viscosity measurements, Josh McCune for assistance with DMA calibration and GEL modulus measurements, and Rebecca Embalabala for helpful discussions regarding cell cycle processes. Biorender was used to create some of the figures in this manuscript.

## Notes

### Competing Interest Statement

The authors have declared no competing interest.

