## Supplemental information for "Substrate stiffness and shear stress collectively regulate the inflammatory phenotype in cultured human brain microvascular endothelial cells"

| Supplementary Table 1: Antibodies used for Immunocytochemistry | | | | |
|---|---|---|---|---|
| Antibody | Vendor | Type | Concentration | Secondary |
| **PECAM-1** | Invitrogen #PA5-143217 | Primary, G | 1:50 | Anti-goat 647, 1:1000 |
| **ICAM-1** | Invitrogen # MHCD5401 | FITC conjugate | 1:200 | N/A |
| **Claudin 5** | Invitrogen #352588 | Conjugated to Alexa 488 | 1:100 | N/A |
| **ZO-1** | Invitrogen #MA3-39100-A488 or A594 | Conjugated to Alexa 488 or 594 | 1:100 | N/A |
| **Phalloidin/F-Actin** | Invitrogen #A12379 (488), Invitrogen # A22287 (647) | Conjugated to Alexa 488 or 647 | 1:400-1:500 (66uM stock) | N/A |

| Supplementary Table 2: DMA analysis of GEL samples and determination of Young's Modulus | | | |
|---|---|---|---|
| **Sample (DMA)** | **Young's Modulus (kPa) DMA** | **Shear Modulus (kPa) rheology** | **Young's Modulus (kPa) rheology** |
| 15% GEL, 0.5% strain/min | 40.7, 46.23, 82.72 | 11.7, 11.27, 8.18 | 33.99, 31.08, 23.73 |
| 6.5% GEL, 0.5% strain/min | 6.46, 4.096, 3.501 | 2.79, 2.79, 2.04, 1.82, 1.64, 1.73 | 6.4 (Avg) kPa |

| Supplementary Table 3: Differentially expressed genes between paired comparisons | | | | |
|---|---|---|---|---|
| Comparison | % genes log2FoldChange > 0 | Down (p < 0.1) | Up (p < 0.1) | Total genes |
| R1: 6.5% vs 15% (1.7 dyne) | 15.1% | 1271, 7.5% | 1301, 7.6% | 17032 |
| R2: 6.5% vs 15% (0 dyne) | 43% | 3823, 22% | 3570, 21% | 17032 |
| R3: 0 vs 1.7 dyne (6.5%) | 52% | 4357, 26% | 4398, 26% | 17032 |
| R4: 0 vs 1.7 dyne (15%) | 47% | 3989, 23% | 4004, 24% | 17032 |

| Supplementary Table 4: Top 10 significant DEGs per paired results comparison | | | | | |
|---|---|---|---|---|---|
| Result 1: 6 kPa vs 30 kPa GEL at 1.7 dyne/cm$^2$ FSS | | | | | |
| | baseMean | log2FoldChange | pvalue | padj | gene_sig1 |
| symbol | | | | | |
| CXCL8 | 11391.6507 | -5.1699841 | 1.26E-83 | 2.10E-79 | down |
| CCL2 | 15228.039 | -2.8471836 | 7.66E-82 | 6.37E-78 | down |
| HDAC9 | 1281.08296 | -3.0944663 | 3.16E-64 | 1.75E-60 | down |
| MAMDC2 | 2276.96614 | 1.49634965 | 7.67E-58 | 3.19E-54 | up |
| CXCL1 | 7146.69849 | -2.7070704 | 1.39E-36 | 4.62E-33 | down |
| IL1A | 534.458077 | -3.0349563 | 8.07E-34 | 2.24E-30 | down |
| MPZL2 | 2739.4506 | 1.45505609 | 8.15E-31 | 1.94E-27 | up |
| ICAM1 | 17533.8426 | -2.8810025 | 3.70E-28 | 7.68E-25 | down |
| COL8A1 | 15426.9681 | 0.75422676 | 2.11E-27 | 3.91E-24 | ns |
| RGS5 | 52338.6464 | 0.97938075 | 2.53E-26 | 4.21E-23 | ns |
| Result 2: 6 kPa vs 30 kPa GEL at 0 dyne/cm$^2$ FSS | | | | | |
| symbol | baseMean | log2FoldChange | pvalue | padj | gene_sig2 |
| UBD | 971.547915 | -6.8314964 | 4.98E-208 | 8.45E-204 | down |
| IL1RL1 | 28365.1999 | -2.4916998 | 5.84E-154 | 4.96E-150 | down |
| MMP10 | 3144.6494 | -4.893583 | 1.14E-149 | 6.46E-146 | down |
| TNFSF15 | 15569.5944 | -4.3508559 | 7.96E-149 | 3.37E-145 | down |
| DCBLD2 | 11595.1442 | -2.4272081 | 2.19E-148 | 7.42E-145 | down |
| COL8A1 | 15426.9681 | -1.7398816 | 1.99E-141 | 5.62E-138 | down |
| CXCL8 | 11391.6507 | -6.1762567 | 4.95E-136 | 1.20E-132 | down |
| CXCL1 | 7146.69849 | -4.4477545 | 6.24E-123 | 1.32E-119 | down |
| PPL | 286.572506 | 4.77686404 | 2.04E-122 | 3.85E-119 | up |
| PVR | 14287.375 | -1.6557224 | 2.65E-122 | 4.50E-119 | down |
| Result 3: 0 vs 1.7 dyne/cm$^2$ FSS for 6 kPa GEL | | | | | |
| symbol | baseMean | log2FoldChange | pvalue | padj | gene_sig3 |
| LIPG | 21325.6626 | -3.7500367 | 0 | 0 | down |
| IL1RL1 | 28365.1999 | -4.2882538 | 0 | 0 | down |
| CCL2 | 15228.039 | 5.14049572 | 4.43E-269 | 2.50E-265 | up |
| LAP3 | 17008.5838 | 2.1566841 | 9.42E-225 | 4.00E-221 | up |
| BPGM | 6794.45798 | -2.3329793 | 8.63E-217 | 2.93E-213 | down |
| STC2 | 9091.22376 | -3.4219127 | 2.03E-212 | 5.74E-209 | down |
| SRPX2 | 18091.8675 | -2.840914 | 1.61E-183 | 3.91E-180 | down |
| INSYN2B | 3057.19558 | -2.1803373 | 5.66E-182 | 1.20E-178 | down |
| NOL4L | 9844.23535 | -1.4357894 | 2.91E-162 | 5.49E-159 | down |
| RELN | 2915.31925 | -4.9143014 | 8.64E-160 | 1.47E-156 | down |
| Result 4: 0 vs 1.7 dyne/cm$^2$ FSS for 30 kPa GEL | | | | | |

| symbol | baseMean | log2FoldChange | pvalue | padj | gene_sig4 |
|---|---|---|---|---|---|
| CCL2 | 15228.039 | 4.71064164 | 8.64E-265 | 1.47E-260 | up |
| GBP4 | 1716.95321 | 6.0141775 | 3.36E-238 | 2.85E-234 | up |
| MAMDC2 | 2276.96614 | 2.83381319 | 6.79E-213 | 3.84E-209 | up |
| DUSP4 | 13817.5045 | -2.0723435 | 7.56E-207 | 3.21E-203 | down |
| FRMD6 | 20716.1083 | 1.45479783 | 1.17E-179 | 3.97E-176 | up |
| TMEM37 | 1169.42652 | -2.9456966 | 1.92E-174 | 5.44E-171 | down |
| CXCL1 | 7146.69849 | 5.2124995 | 2.27E-162 | 5.49E-159 | up |
| LAP3 | 17008.5838 | 1.78805194 | 1.19E-153 | 2.53E-150 | up |
| SAMHD1 | 8225.16981 | 2.46101629 | 1.35E-147 | 2.55E-144 | up |
| IFIT1 | 1137.49821 | 6.25403485 | 1.64E-144 | 2.79E-141 | up |

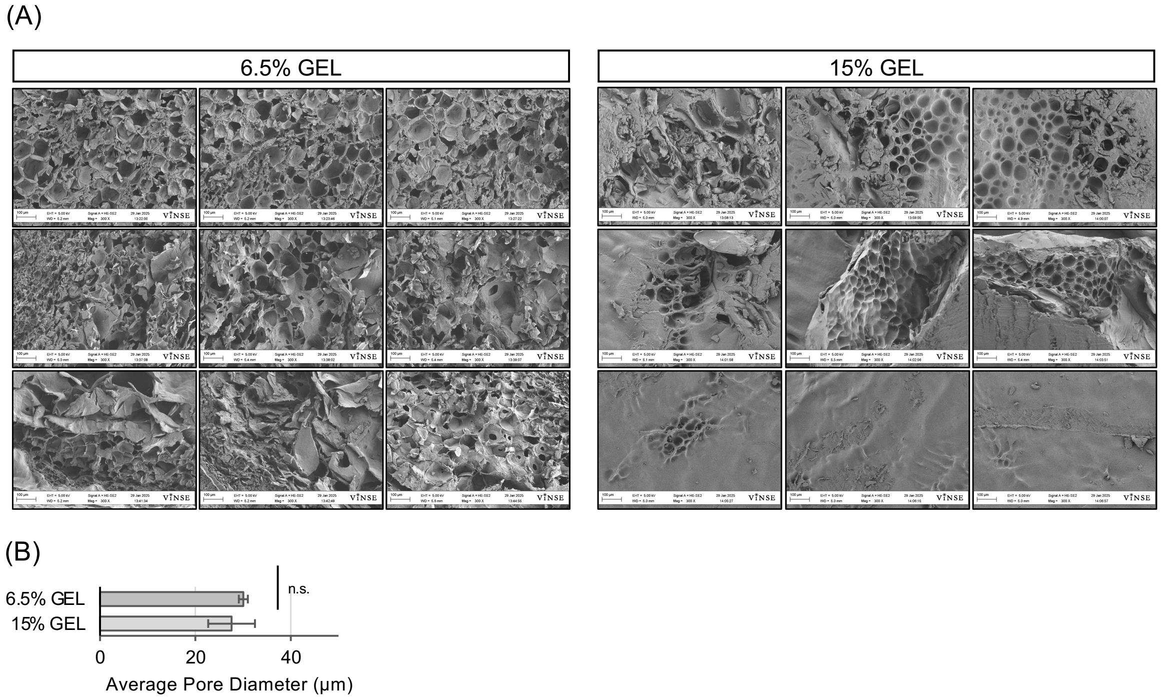


**Supplemental Figure 1: Analysis of GEL pore size with SEM.**
(A) Representative SEM images of 6.5% GEL (6 kPa) and 15% GEL (30 kPa) Three samples of each GEL were measured (each row represents a single sample) and three images were taken per sample. (B) Student's t-test was used to determine significance between dehydrated pore size. No significant difference in pore size was seen between GEL samples. Data are presented as mean ± SD.

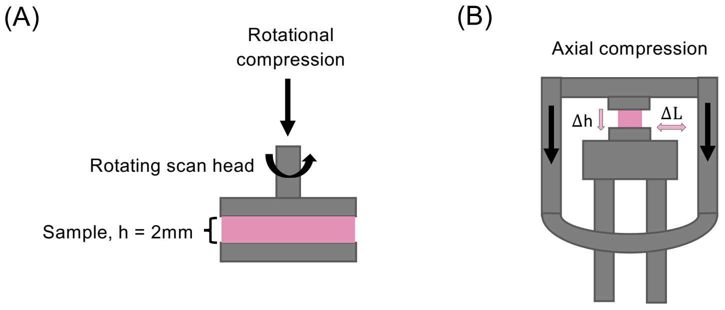


$$(1)\ E^{*} = 2G^{*}(1+v)$$

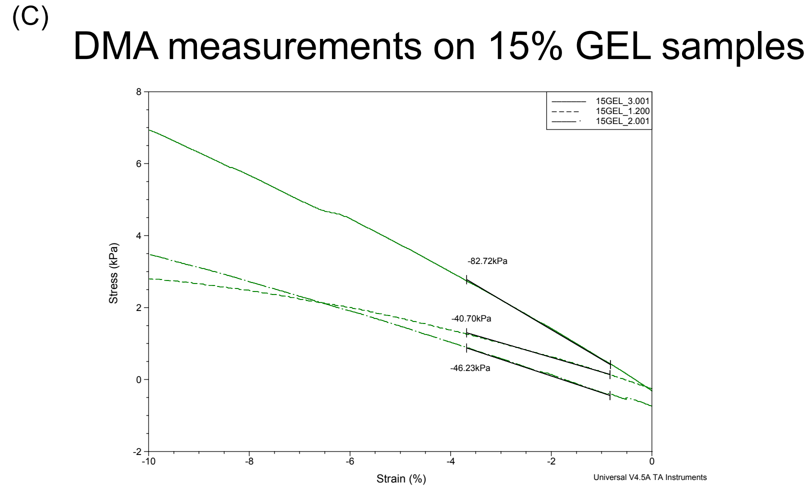


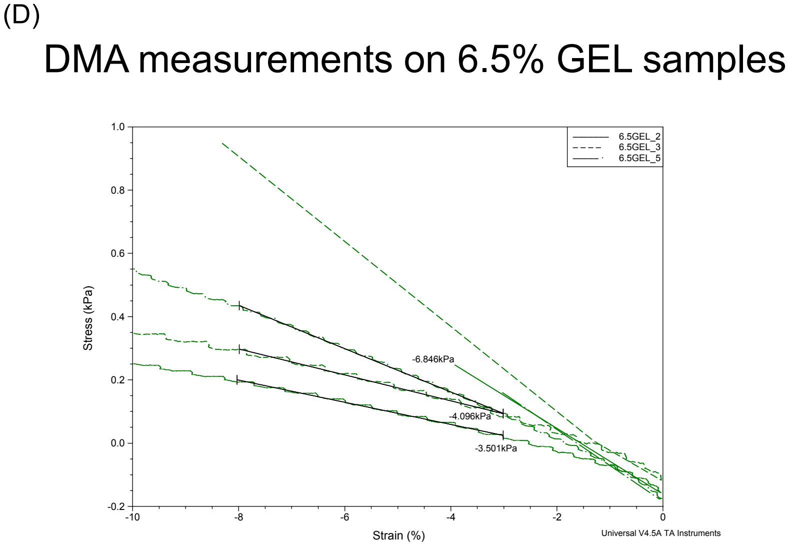


**Supplementary Figure 2: Dynamic mechanical analysis of hydrated GEL samples.**
(A) Representation of parallel plate rheology for measurements of GEL shear modulus and media viscosity. (B) Representation of axial compression set up for dynamic mechanical analysis of GEL samples for direct measurement of the Young's modulus. Equation 1 describes conversion between complex Young's Modulus (E*) and complex shear modulus (G*) (C) DMA measurements of Young's

modulus in compressive stress were collected on n=3 replicates of 15% GEL samples. (D) DMA measurements in compressive stress were repeated for n=3 samples of 6.5% GEL. Table 2 shows Young's modulus measurements in addition to the conversion of rheological measurements of the shear modulus to the Young's modulus.

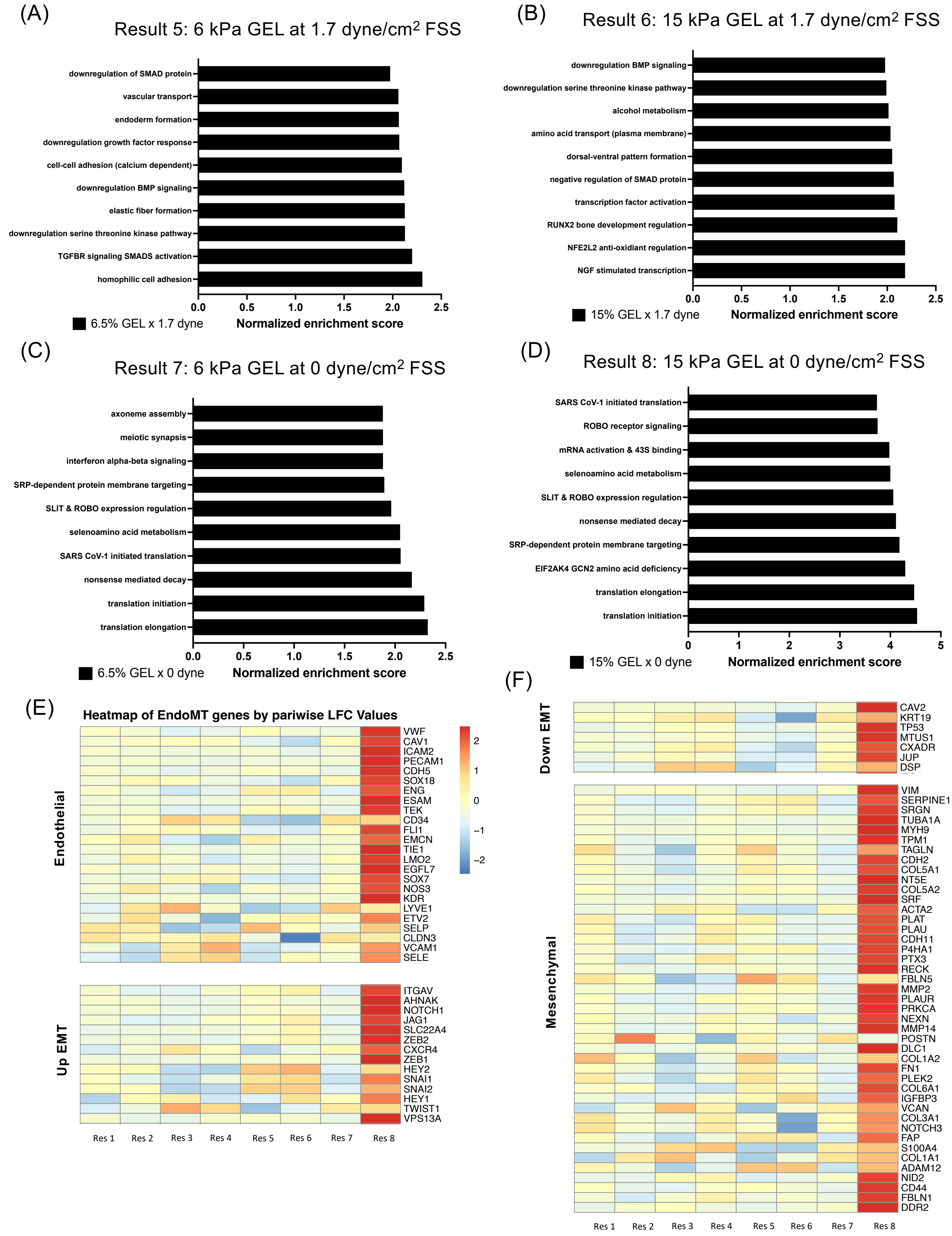


**Supplementary Figure 3: Pathway analysis of 1 vs all comparisons and EndoMT signature genes.** (A-D) Similar to Figure 4 in the main text, graphs display top 10 upregulated pathways from pathway analysis (ranked by normalized enrichment score) when comparing a single condition versus a group of all remaining conditions. (E-F) Heatmap of log2foldchange across paired results and Result 1 versus all results for signature genes related to the endothelial-to-mesenchymal transition (EndoMT) that is a marker for endothelial dysfunction. Result 1 refers to 6 kPa vs 30 kPa GEL at 1.7 dyne/cm$^2$ FSS, Result 2 refers to 6 kPa vs 30 kPa GEL at 0 dyne/cm$^2$ FSS, Result 3 refers to 0 vs 1.7 dyne/cm$^2$ FSS for 6 kPa GEL, and Result 4 refers to 0 vs 1.7 dyne/cm$^2$ FSS for 30 kPa GEL, Result 5 refers to 6 kPa at 1.7 dyne/cm$^2$ vs all conditions, Result 6 refers to 30 kPa at 1.7 dyne/cm$^2$ vs all conditions, Result 7 refers to 6 kPa at 0 dyne/cm$^2$ vs all conditions, and Result 8 refers to 30 kPa at 0 dyne/cm$^2$ vs all conditions. EndoMT-related genes were selected from Evrard SM, et al. (2016).[1]

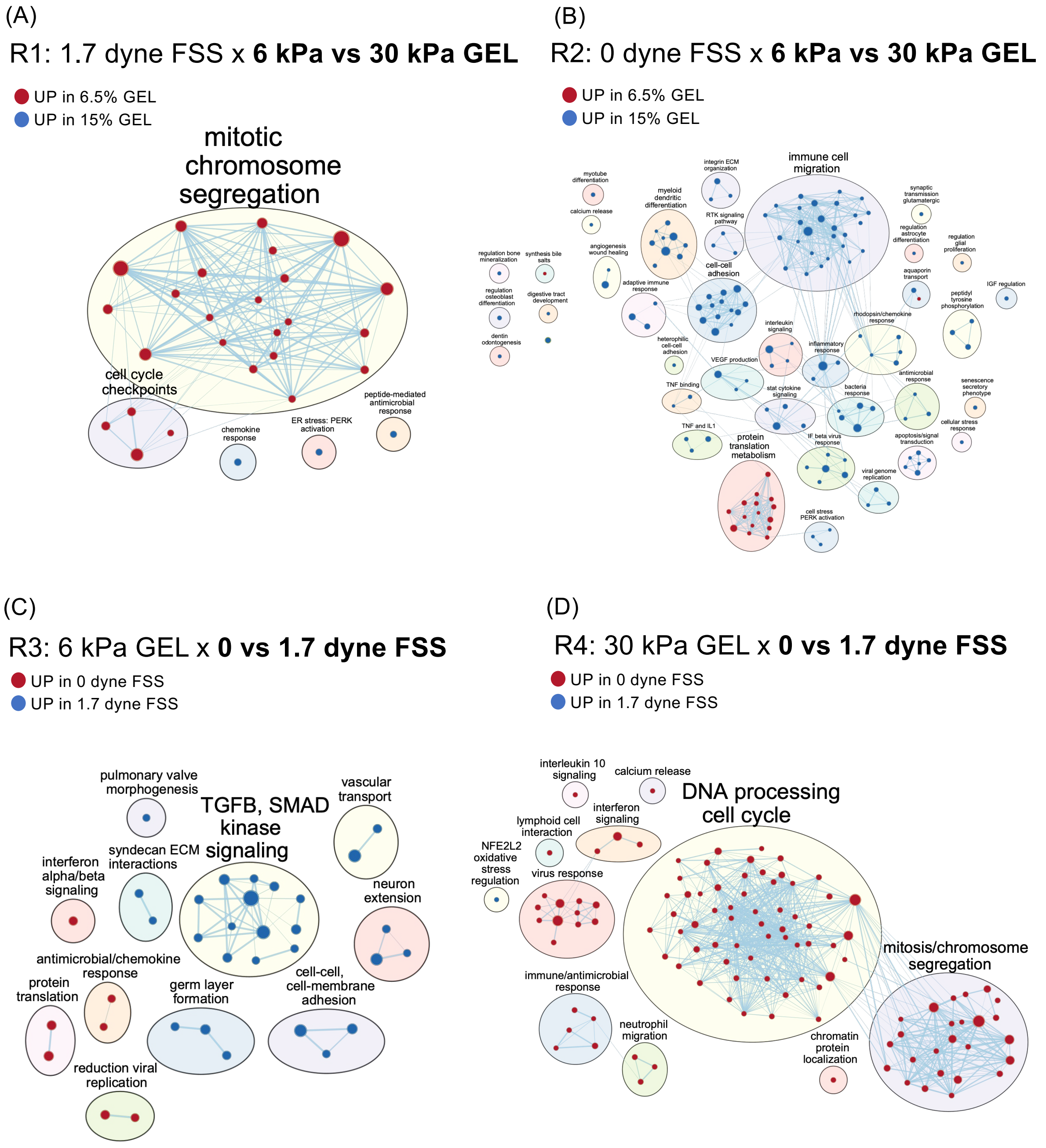


**Supplementary Figure 4: Enrichment maps of significant cellular pathways across paired comparisons.**

As described in Methods, enrichment maps were generated using cut-off values of $p < 0.005$ and FDR q-value < 0.01 from GSEA results. Shown are all significant nodes, representing gene sets from Reactome, Kegg Medicus, and Gene Ontology databases. Cluster labels were manually curated from initial WordCloud app in Cytoscape and gene set names.

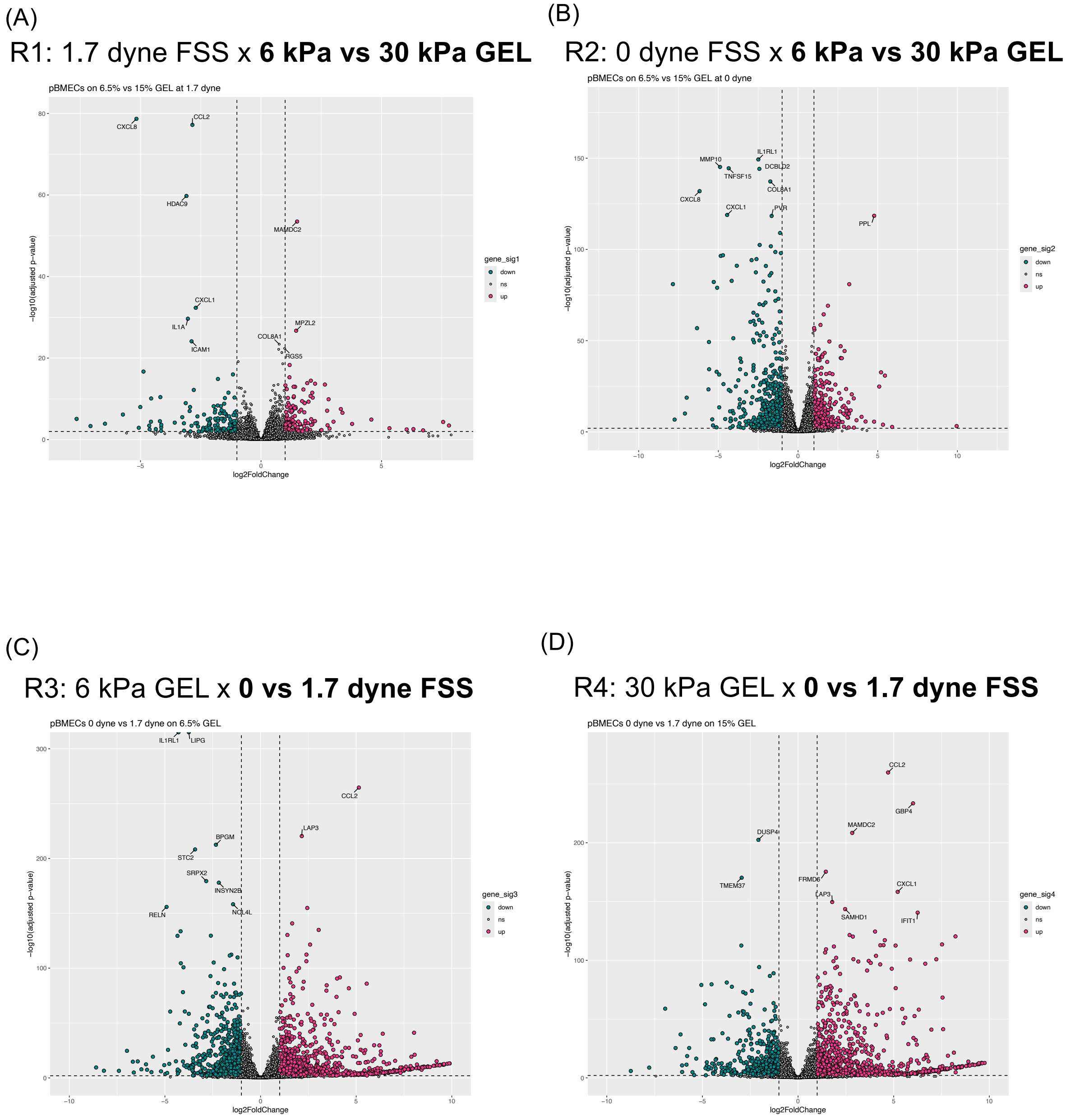


**Supplemental Figure 5: Top 10 DEGs between paired comparisons ranked by adjusted p-value.** Volcano plots of paired comparisons between experimental conditions. Significance cutoffs (dashed lines) represent adjusted p-value (padj) < 0.01 and log2fold change ± 1. Annotated genes represent top 10 genes ranked by adjusted p-value. Genes colored pink indicating significant upregulation in the first term of the comparison, or turquoise, indicating upregulation in the second term of the comparison. Genes colored in grey have padj < 0.1 but log2foldchange between -1 and 1. See Supplemental Table 3 for the log2foldchange and p-adj values for all annotated genes.

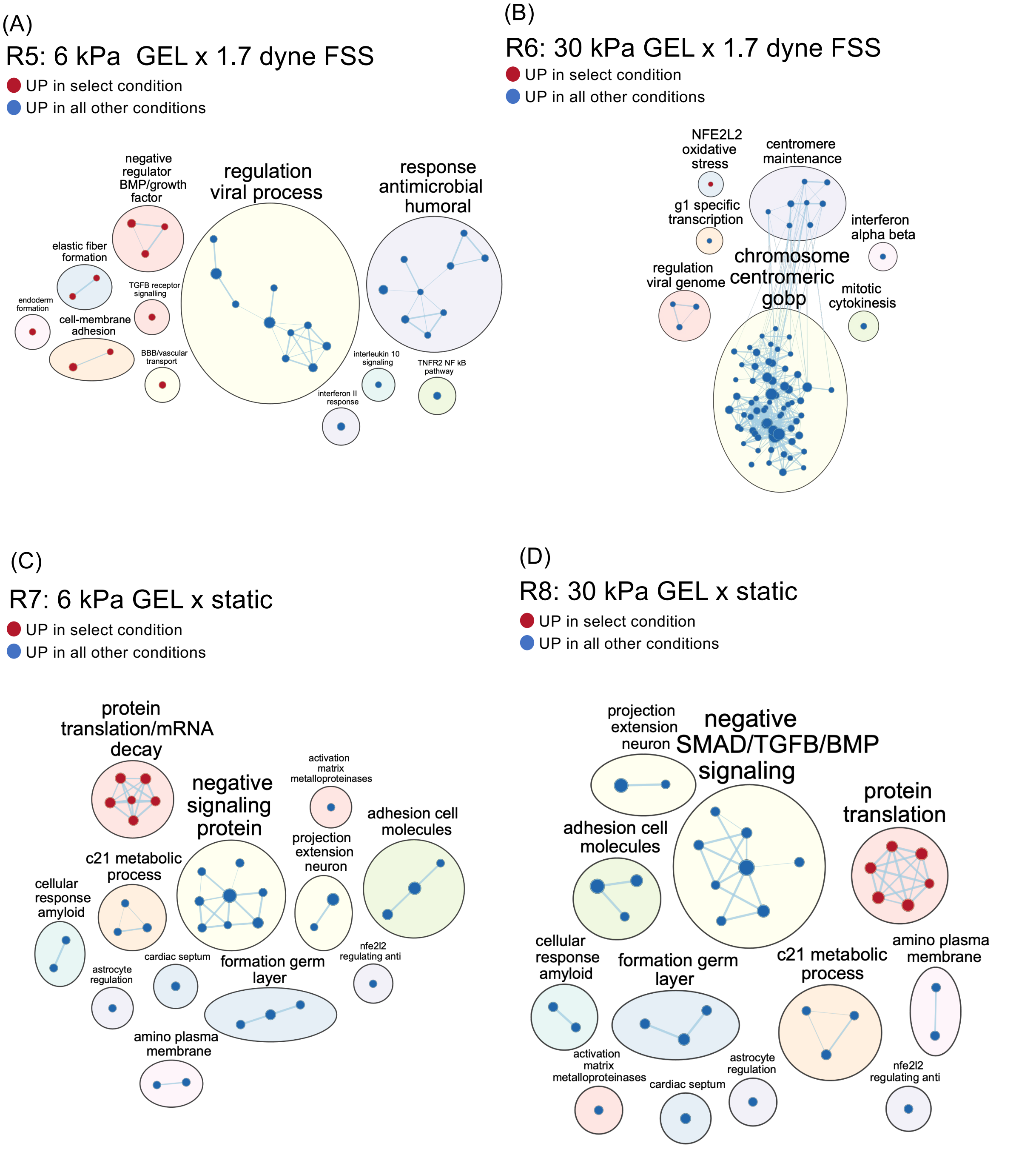


**Supplemental Figure 6: Enrichment maps from pathway analysis of 1 condition versus all other conditions.**
As described in Methods, enrichment maps were generated using cut-off values of $p < 0.005$ and FDR q-value < 0.01 from GSEA results. Shown are all significant nodes, representing gene sets from Reactome, Kegg Medicus, and Gene Ontology databases. Cluster labels were manually curated from initial WordCloud app in Cytoscape and gene set names.

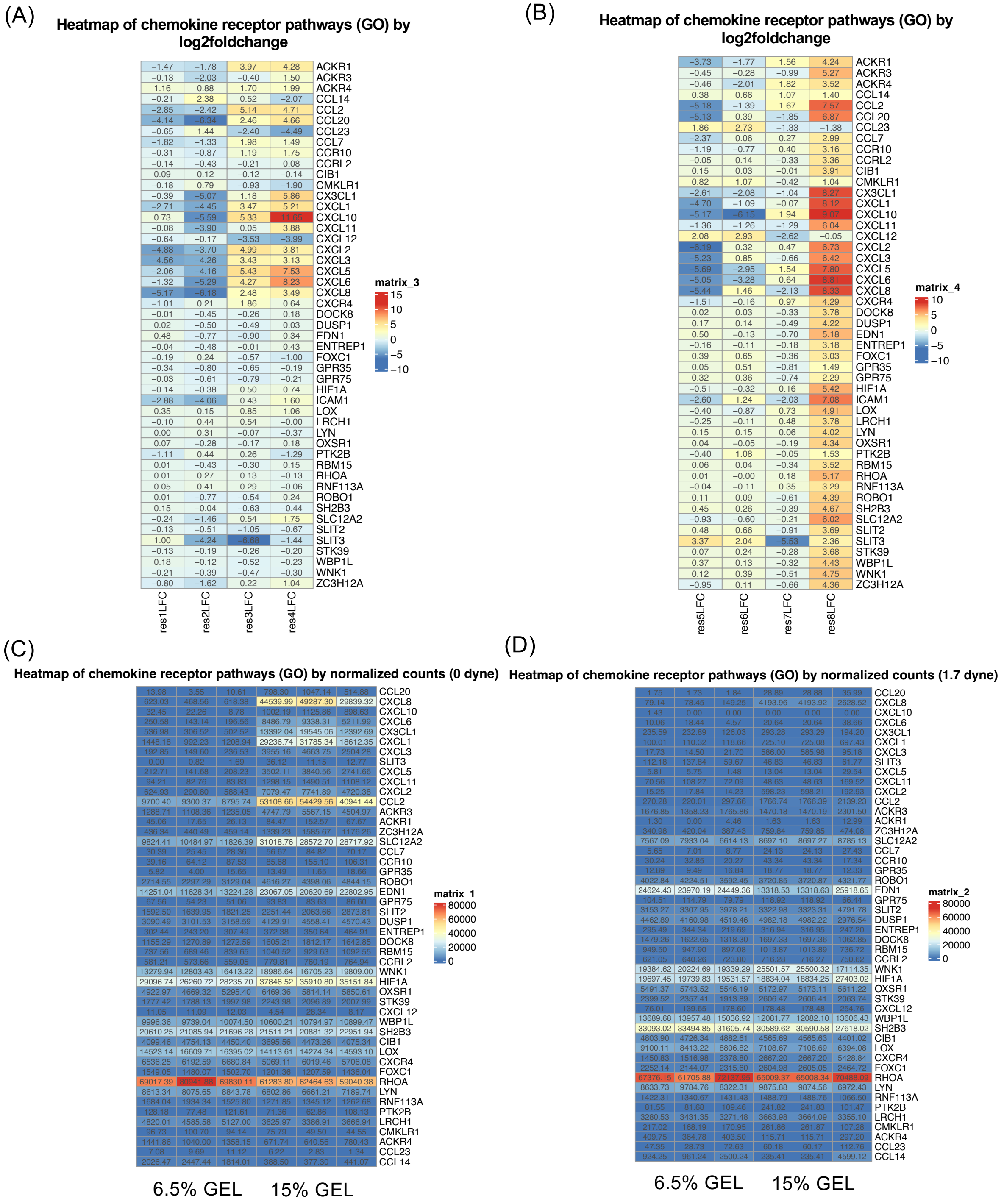


**Supplemental Figure 7: Heatmap of log2foldchange and count data for Chemokine Receptor Pathway genes.**

Using the Chemokine Receptor Pathway gene set (Gene Ontology), log2foldchange values are displayed as a heatmap for results 1-4 (panel A) and results 5-8 (panel B). Similarly, count data is displayed in a heatmap for all samples under 0 dyne/cm$^2$ FSS (panel C) and 1.7 dyne/cm$^2$ FSS (panel D). Three samples were analyzed per condition and are grouped by GEL (see bottom of heatmap). Result 1 refers to 6 kPa vs 30 kPa GEL at 1.7 dyne/cm$^2$ FSS, Result 2 refers to 6 kPa vs 30 kPa GEL at 0 dyne/cm$^2$ FSS, Result 3 refers to 0 vs 1.7 dyne/cm$^2$ FSS for 6 kPa GEL, Result 4 refers to 0 vs 1.7 dyne/cm$^2$ FSS for 30 kPa GEL,

Res 5ult refers to 6 kPa at 1.7 dyne/cm$^2$ vs all conditions, Result 6 refers to 30 kPa at 1.7 dyne/cm$^2$ vs all conditions, Result 7 refers to 6 kPa at 0 dyne/cm$^2$ vs all conditions, and Result 8 refers to 30 kPa at 0 dyne/cm$^2$ vs all conditions.
